# In and out of replication stress: PCNA/RPA1-based dynamics of fork stalling and restart in the same cell

**DOI:** 10.1101/2024.12.03.626292

**Authors:** Teodora Dyankova-Danovska, Sonya Uzunova, Georgi Danovski, Rumen Stamatov, Petar-Bogomil Kanev, Aleksandar Atemin, Aneliya Ivanova, Stoyno Stoynov

## Abstract

Replication forks encounter various impediments which induce fork stalling and threaten genome stability, yet the precise dynamics of fork stalling and restart at the single-cell level remain elusive. Herein, we devise a live-cell microscopy-based approach to follow hydroxyurea-induced fork stalling and subsequent restart at 30-s resolution. We measure two distinct processes during fork stalling. One is rapid PCNA removal, which reflects the drop in DNA synthesis. The other is gradual RPA1 accumulation on up to 2400 nt of ssDNA per fork despite an active intra-S checkpoint. Restoring the nucleotide pool enables prompt restart without post-replicative ssDNA and a smooth cell cycle progression. ATR, but not ATM inhibition, accelerates hydroxyurea-induced RPA1 accumulation nine-fold, leading to RPA1 exhaustion within 20 min. Fork restart under ATR inhibition led to the persistence of ∼600 nt ssDNA per fork after S-phase, which reached 2500 nt under ATR/ATM co-inhibition, with both scenarios leading to mitotic catastrophe. MRE11 inhibition had no effect on PCNA/RPA1 dynamics regardless of ATR activity. E3 ligase RAD18 was recruited at stalled replication forks in parallel to PCNA removal. Our results shed light on fork dynamics during nucleotide depletion and provide a valuable tool for interrogating the effects of replication stress-inducing anti-cancer agents.

## Introduction

Mammalian DNA replication is a gargantuan task that requires the spatiotemporally coordinated stochastic firing of replication origins that number in the tens of thousands [1]. As replication forks traverse the genome, they inevitably encounter various impediments, including DNA lesions (e.g. ssDNA gaps, UV-induced bulky lesions, interstrand crosslinks, and DNA-protein crosslinks), transcription machinery, or inherently difficult-to-replicate regions [2]. In addition to such “physical” barriers, oncogene stimulation and nucleotide depletion may also limit replication fork progression [3]. All of the above-described can act as triggers of replication stress, a state whereby conventional fork progression and origin firing are dysregulated, leading to lower replication fidelity and DNA breakage [4]. While a number of DNA damage response mechanisms can resolve or tolerate impediments to replication fork progression, beyond a certain threshold of damage or nucleotide depletion, forks can collapse and give rise to an excess of double-strand breaks (DSBs), leading to cell death. Enhanced mitogenic signaling and DNA repair defects, which are commonly observed in tumors, elicit a chronic state of tolerable replication stress, which is considered a major driver of genomic instability and tumor evolution [5]. “Pushing” cancer cells beyond these tolerable levels of replication stress has thus emerged as a therapeutic strategy [6].

To effectively cope with replication stress, eukaryotic cells have evolved an elaborate network of factors that sense and respond to replication stress. This response is directed by the ataxia telangiectasia and Rad3-related (ATR) serine/threonine kinase, which is activated by the generation of ssDNA stretches generated as a consequence of multiple replication stress-associated events [7]. Through the phosphorylation of various substrates, among which replisome components, DNA repair, and checkpoint factors, ATR regulates origin firing, replication fork stability, and cell cycle progression. The essentiality of the ATR-mediated replication stress response is highlighted by the embryonic lethality observed under ATR deficiency [8,9] as well as the severe developmental disorder Seckel syndrome, caused by ATR hypomorphism [10].

Aberrant oncogene activation can induce a depletion of the cellular nucleotide pool, eliciting replication stress and genome instability [11]. Nucleotide depletion by ribonucleotide reductase inhibitor hydroxyurea (HU) leads to the generation of ssDNA that activates the intra-S phase checkpoint-mediated stress response [12,13]. In addition, mild hydroxyurea treatment has been shown to elicit considerable fork reversal (∼20% of forks) and post-replicative ssDNA gaps (∼30% of forks), as visualized by electron microscopy [14]. Inhibition of the intra-S checkpoint causes the uncoupling of DNA unwinding and synthesis [15–17], resulting in the rapid exhaustion of the cellular RPA pool [18].

DNA fiber analyses have revealed an increasing number of factors implicated in fork reversal, degradation, and restart, highlighting protective roles for BRCA1/2 against the nucleolytic degradation of stalled replication forks [19–22]. Further, the proteome of HU-stressed forks has been comprehensively characterized via iPOND coupled with mass spectrometry [23], revealing a wide range of proteins involved in replisome stabilization and the prevention of fork collapse [24], with similar resources generated for camptothecin- and aphidicolin-induced replication stress [25,26]. A number of sequencing-based approaches have been developed to meticulously quantify replication timing, speed, and sites of fork collapse [27–29]. While the precise kinetics of recruitment and dissociation to micro-IR-induced lesions have been characterized for a number of replisome components, including PCNA and RPA1[30,31], replisome dynamics through the entire process of replication fork stalling, reversal/protection, and restart, have not been followed.

Herein, we introduce a live-cell imaging-based approach for interrogating the dynamics of replisome components during replication stress and subsequent recovery at 30-s resolution. We visualized PCNA removal and RPA1 recruitment at stalled replication forks following HU-induced nucleotide depletion, precisely measuring the kinetics of both processes. Thereafter, we followed PCNA recovery and RPA removal during replication fork restart within the same replication foci. The influence of ATR, ATM, and MRE11 activities on fork dynamics during fork stalling and restart were also quantified. Strikingly, our methodology revealed that PCNA dissolution occurs with a half-time of 2 min, while RPA1 recruitment occurs gradually over 1.5 h. In contrast, PCNA recruitment and RPA removal both occurred within minutes during fork recovery. ATR inhibition greatly increased both the rate and extent of RPA1 recruitment, while having no effect on PCNA nor on RPA1 kinetics during fork recovery. Interestingly, 20% of replication stress-induced RPA1 persisted after fork recovery. While ATM inhibition alone had no effect on any of the above-described phenomena, co-treatment with ATRi increased this fraction to 60%, suggesting a protective role for ATM during fork resolution under suppressed ATR signaling.

## Results

### Changes in PCNA and RPA1 distribution reflect replication fork stalling and restart

To follow replication fork dynamics during replication stress and subsequent recovery, we opted for the concurrent visualization of fluorescently tagged PCNA and RPA1 in live cells. As the eukaryotic clamp, PCNA is abundant at the replication fork, with one molecule accommodating leading-strand synthesis and many molecules acting at Okazaki fragments on the lagging strand [32]. It enhances polymerase processivity and serves as a platform for the recruitment of functionally diverse factors at the fork. The amount of PCNA on a fork at a given point in time is determined by the rate of its loading, which reflects replication fork speed, as well as by the rate of its removal after the completion of nascent strand synthesis. Meanwhile, RPA binds single-stranded DNA generated at the replication fork when DNA synthesis at the leading or lagging strand is slower than replicative DNA unwinding [33–35]. Another source of ssDNA is resection by various nucleases during the repair of DNA lesions at the replication fork [36]. Thus, the amount of RPA1 reflects ssDNA generated during unperturbed replication as well as during replication fork stalling and restart.

Herein, we employed HeLa Kyoto cells co-expressing RPA1-EGFP/mouse PCNA-mCherry [37] generated through bacterial artificial chromosome (BAC) recombineering [38]. This line expresses the tagged proteins at near-endogenous levels under the control of their own regulatory elements [39]. Single clone selection was performed to ensure homogenous expression of both proteins across the cell population. Western blot revealed that RPA1-EGFP (hereafter referred to as RPA1) levels were 1:3 those of endogenous RPA1, while mouse PCNA-mCherry (hereafter referred to as PCNA) levels were approximately 1:4.2 those of endogenous PCNA (Figure 1A). To obtain high-resolution time-lapse images, we initially opted for airyscan super-resolution microscopy at 2-min temporal resolution (Figure 1B, C).

**Figure 1.**
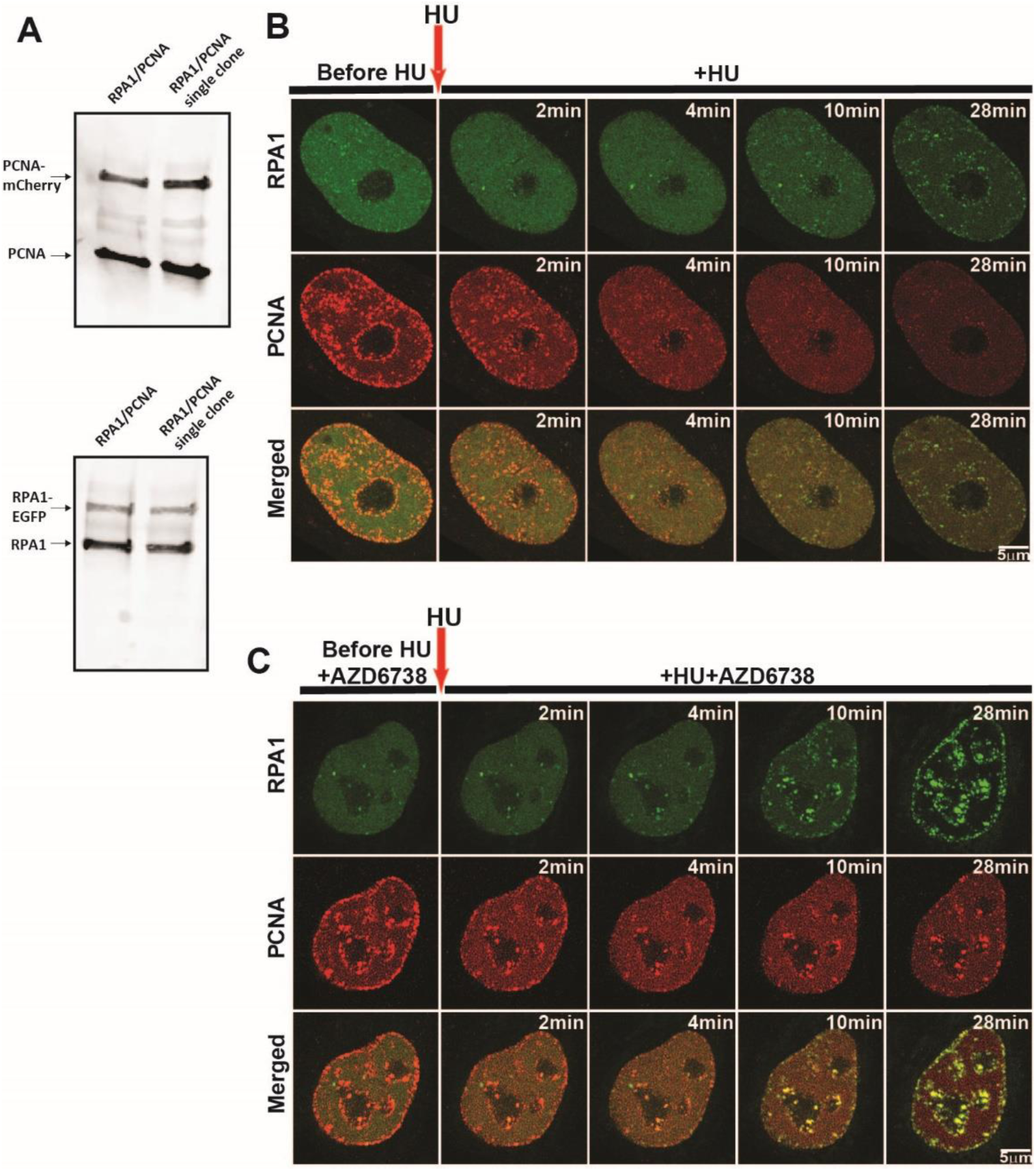
Nuclear distribution of PCNA and RPA1 during hydroxyurea-induced replication stress. (**a**) Comparison of the expression level of BAC-tagged PCNA-mCherry and RPA1-EGFP versus their endogenous counterparts via western blotting. (**b**) Representative timelapse airyscan images of RPA1-EGFP and PCNA-mCherry before and during HU-induced replication stress. Scale bar = 5 µm. (**c**) Same as (b), but in the presence of ATR inhibitor AZD6738 before and during HU treatment. Scale bar = 5 µm. **Abbreviations**: HU: hydroxyurea; AZD: AZD6738.

Addition of 10mM HU led to a rapid dissociation of PCNA foci, which was followed by RPA1 accumulation within the same regions (Figure 1B, Video 1). Inhibiting ATR using AZD6738 [40] led to considerably larger RPA1 foci (Figure 1C, Video 2). However, time-lapse imaging at 2-min was not sufficient to precisely follow the kinetics of PCNA removal and led to substantial photobleaching, which precluded us from studying subsequent fork restart within the same cells. Thus, we opted for spinning-disk microscopy, which enabled us to follow fork stalling and restart in the same cells, at 30-s resolution.

## PCNA and RPA1 dynamics during replication fork stalling and restart

To study PCNA and RPA1 dynamics during replication fork stalling, we imaged cells for 15 min, whereafter we added HU and continued imaging for 1 h. To follow fork restart, we then washed out HU and imaged cells for another 45 min. As observed through airyscan microscopy, nucleotide depletion by HU led to rapid dissolution of mouse PCNA foci, followed by a gradual accumulation of RPA1 within the same nuclear regions (Figure 2A, Video 3).

**Figure 2.**
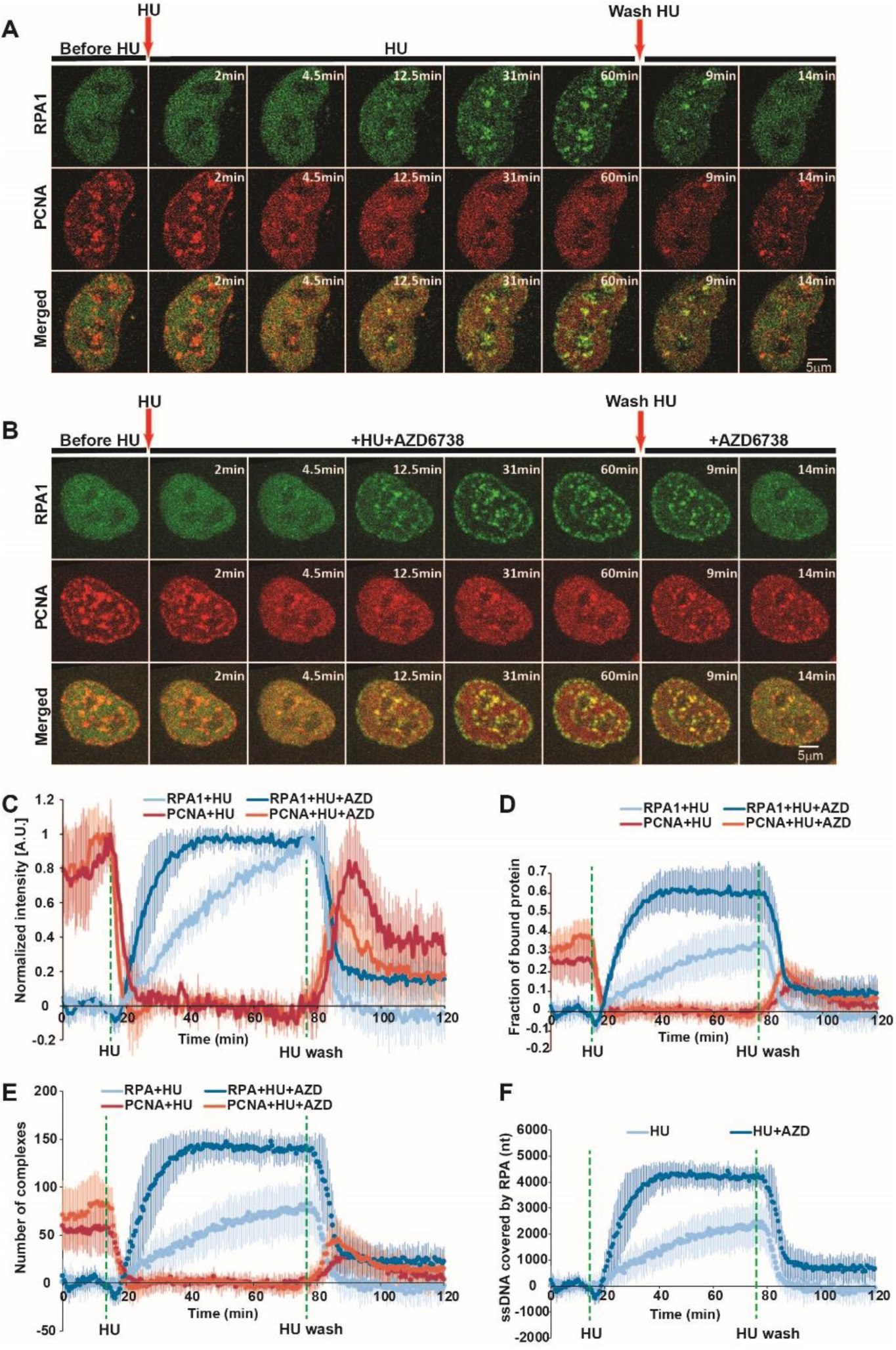
Dynamics of RPA1 and PCNA during HU-induced replication fork stalling and restart. (**a**) Representative time-lapse images of RPA1 and PCNA before, during, and after HU treatment. Arrows indicate timepoints of HU addition and washout. Scale bar = 5 µm. (**b**) Same as (a), but with inhibition of ATR (3µM AZD6738) throughout the experimental period. (**c**) Normalized kinetics of PCNA and RPA1 at replication foci during HU-induced replication fork stalling and restart, with or without ATR inhibition. The maximum intensity of PCNA/RPA at replication foci is normalized to 1. (**d**) Fraction of PCNA and RPA1 bound at replication foci during HU-induced replication fork stalling and restart, with or without ATR inhibition, relative to the total nuclear intensity of PCNA/RPA1, which is normalized to 1. (**e**) Estimated number of PCNA homotrimer and RPA heterotrimer complexes engaged at replication foci with or without ATR inhibition. (**f**) Estimated number of nucleotides covered by RPA heterotrimers with or without ATR inhibition. For HU only: n = 17 cells; for HU+AZD: n = 10 cells. **Abbreviations**: HU: hydroxyurea; AZD: AZD6738.

To determine whether there are differences in the dynamics of mouse and human PCNA, we employed a double BAC-tagged cell line harboring both proteins of interest (mouse PCNA-mCherry/human PCNA-EGFP). The two homologues exhibited identical kinetic behavior under HU treatment (Figure S1), justifying the use of mouse PCNA-mCherry for our subsequent experiments.

While single foci can be tracked and measured, the variation of their size and shape complicates the accurate measurement of the total amount of a given protein engaged at replication foci throughout the nucleus. Thus, we opted for measuring the changes in freely diffusing PCNA/RPA1 at nuclear sites without visible foci during replication fork stalling and restart (Figure 3A). The dissolution of PCNA foci after HU addition led to a considerable increase in intensity at these sites, suggesting that the removed PCNA indeed becomes part of the freely diffusing pool (Figure 2C and 3). Therefore, following the fluctuations in freely diffusing PCNA at these regions enabled us to measure the amount of PCNA that is unloaded from replication forks (Figure 3C-E).

**Figure 3.**
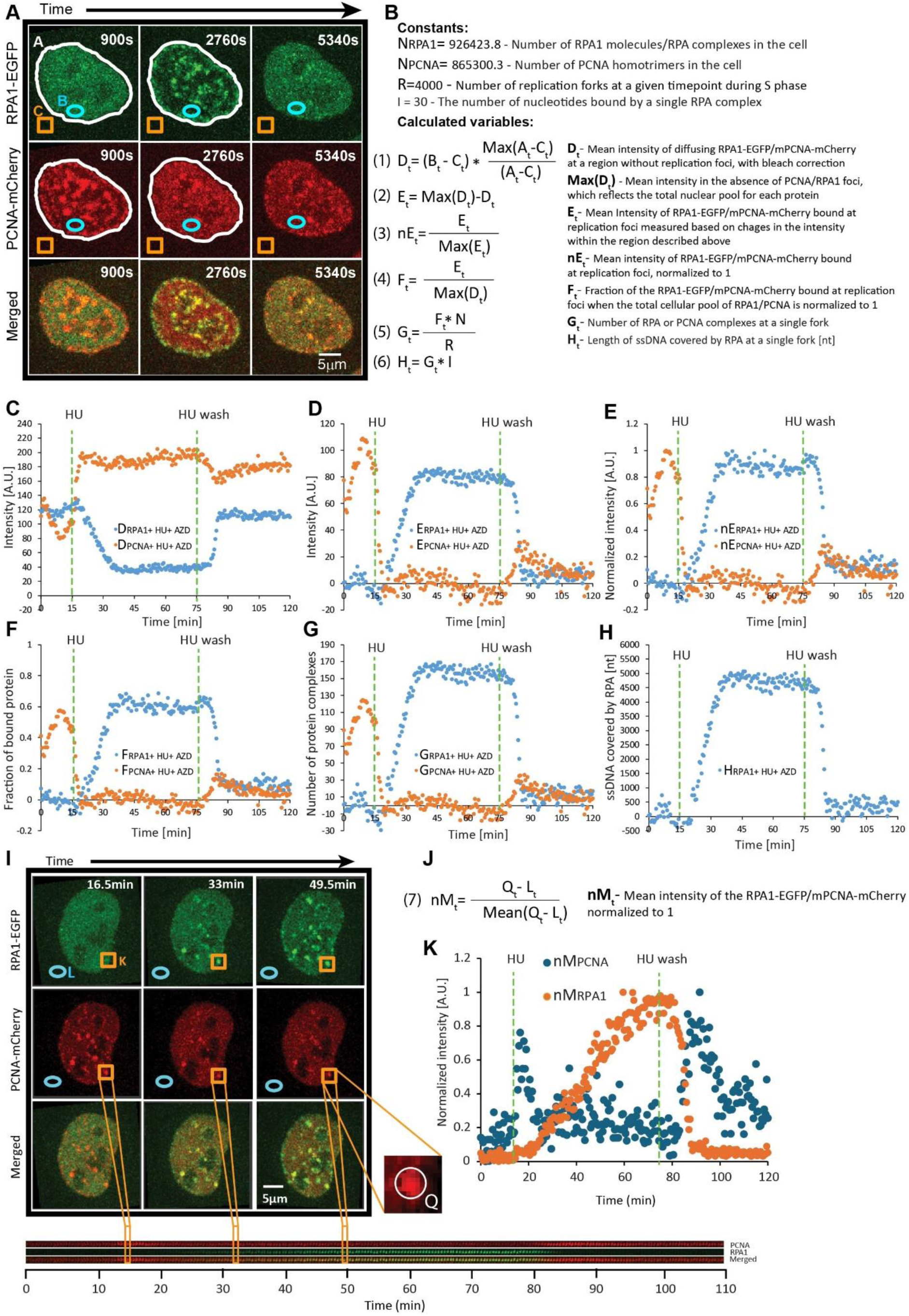
Detailed overview for the measurement and quantification of PCNA and RPA1 engaged at replication foci during fork stalling and restart. (**a**) Regions of interest applied for analysis. ‘A’ represents the cell nucleus (white lining), ‘B’ represents a region within the nucleus without replication foci (blue ellipse), and ‘C’ represents noise outside of cells (orange square). (**b**) Constants and formulas for the calculation of variables used for quantifying RPA1 and PCNA kinetics. (**c**) Mean intensity of diffusing RPA1-EGFP/mPCNA-mCherry within region ‘B’, calculated as shown in the formula for D_t_. (**d**) Mean intensity of RPA1-EGFP/mPCNA-mCherry bound at replication foci, calculated as per the formula for E_t_ (2). (**e**) Same as (d), but with the maximum intensity normalized to 1, calculated as per the formula for nE_t_ (3). (**f**) Fraction of the RPA1-EGFP/mPCNA-mCherry bound at replication foci when the total cellular pool of RPA1/PCNA is normalized to 1, as per formula (4). (**g**) Estimated number of RPA and PCNA complexes engaged at a single replication fork, calculated as per the equation for G_t_ (5). (**h**) Average length of ssDNA (nt) covered by RPA at a replication fork, calculated as per the formula for H_t_ (6). (**i**) Representative timelapse images of single RPA1/PCNA foci tracking. After a focus is tracked, a square region (orange square) surrounding is cropped, and a kymogram is created (below). The background noise is measured in a region indicated by the blue circle. (**j**) Formula (7) for calculating the mean intensity of the RPA1-EGFP/mPCNA-mCherry signal in a single focus (Q_t_) normalized to 1. (**k**) Normalized intensity of the single tracked focus (nM_t_) from (i).

To determine what fraction of the total PCNA is engaged at replication foci, we first measure the mean intensity within the ROIs without visible foci after HU treatment (after the foci were dissolved) (Figure 2D, Figure 3B, F). As the foci have dissolved, the mean intensity within ROIs at this time point reflects that total nuclear pool. Thus, subtracting intensities at the same ROI during stalling and restart from this mean intensity (representing the total fraction) yields an intensity value that reflects the protein bound at foci. Finally, dividing this derived intensity of the bound protein by the intensity reflecting the total pool yields the fraction of protein bound at foci (Figure 3). We employ the same approach to measure mean RPA1-EGFP intensity within the same ROIs, with the difference being that we use the intensity values prior to HU addition to measure the total nuclear pool (because there are no visible RPA1 foci before HU treatment). It should be noted that not all PCNA foci fully dissolve and some RPA1 is engaged at forks even if not visible. However, these contribute negligible systematic error, as shown in our measurements for the dissociation of large PCNA foci (see below).

Taking advantage of the above-described analytical pipeline, we set out to quantify the dynamics of PCNA and RPA1 during HU-induced replication fork stalling and restart at 30-s resolution at the single-cell level. Our measurements demonstrate that within 5 min of HU addition, PCNA foci almost completely dissociated (t_1/2_ = 2.1±0.9 min)(Figure 2C and Supplementary Table 1). In the largest foci, some residual PCNA could still be observed after the complete dissolution of their smaller counterparts. Quantification revealed that approximately 27% of the total nuclear PCNA was engaged at replication foci prior to HU treatment. Interestingly, HU-induced RPA1 accumulation occurred with a much slower half-time (t_1/2_ = 23.9±2.1 min)(Figure 2C and Supplementary Table 1). At the end of the 1-h HU treatment period, 35% of nuclear RPA1 was engaged at replication foci (Figure 2D). Under 3-h HU treatment, this fraction increased to 40%, plateauing after 90 min (Figure S2, Video 4). As early as 5 min after HU washout, RPA1 began to rapidly dissociate from restarting forks, with complete removal observed approx. 12 min after washout. While no RPA1 foci persisted during fork restart after 1-h HU treatment (Figure 2D), 4-5% of the RPA1 engaged at foci remained after wash-out in the 3-h HU treatment condition (Figure S2). This indicates that the ssDNA generated during 1-h nucleotide depletion can be promptly resolved during fork restart, while prolonged depletion gives rise to the persistence of some RPA-coated ssDNA regions. Thus, we selected the 1-h HU treatment conditions for our subsequent experiments.

Upon HU wash-out, PCNA re-accumulation ensued at the previously dissolved replication foci (Figure 2D), reaching a peak within approximately 15 min (t_1/2_ = 5.1±2 min)(Supplementary Table 1). PCNA levels began to decrease shortly thereafter, reflecting S-phase exit. We then sought to determine whether individual PCNA/RPA1 foci exhibit consistent kinetics during HU treatment and washout. To this end, we employed MtrackJ [41] to track and SPARTACUSS [42] to measure individual PCNA/RPA1 foci, which were sufficiently large for tracking (Figure 3I-K). Individual foci exhibited kinetics that closely recapitulate those obtained by measuring the fluctuations in freely diffusing protein (Figure S3A-C).

### Effect of acute nucleotide depletion on cell cycle progression

To determine whether the 1-h treatment with 10mM HU we used for replication stress induction had an effect on cell cycle progression, we followed cells at 10-min intervals for 20 h after HU washout (Figure S4A,D). For 20 h, all cells successfully exited S phase and progressed through G2, mitosis, G1, and into a second S phase. Thereafter, 30% of the cells reached a second mitosis, without unusual RPA1 accumulation at any point during the cycle. This indicated that cells effectively recovered from HU-induced replication fork stalling and proceeded through multiple divisions without harboring unresolved DNA damage. Taken together, our measurements reveal that despite active ATR signaling, RPA continuously accumulates at stalled forks, albeit at a much slower rate than that of PCNA removal. Meanwhile, PCNA-re-accumulation and RPA removal occur at comparable rates during fork restart, which is followed by smooth progression through multiple cell cycles.

### Quantification of PCNA and RPA complexes engaged at forks during stalling and restart

As discussed above, our fluorescence intensity-based data enable us to determine the fraction of PCNA/RPA1 engaged at all replication forks at different timepoints. Combining these with quantitative proteomics data, which are available for the HeLa Kyoto cell line used in this study [39], allows us to infer the number of protein molecules engaged at replisomes during and after replication stress induction. The HeLa proteome contains approximately 2.6 million molecules of PCNA arranged in ∼865,300 homotrimers and 926,424 RPA1 molecules forming the same number of RPA heterotrimer complexes. The total number of forks active at a given timepoint during replication was recently shown to be around 4,000, with the exception of the very early and late stages of S-phase [29]. Dividing the number of protein molecules engaged at all forks by the total number of forks could therefore give us an estimate of how many molecules are present at a single fork (Supplementary Table 2). Thus, we estimated that around 59±22.6 PCNA homotrimers were present at single forks prior to HU addition. To determine the amount of residual PCNA following nucleotide depletion, we selected the largest PCNA foci, as their signal intensity could still be measured after the HU-induced decrease. Our measurements demonstrated that 17.6±6.7% of PCNA remain per replication fork after HU treatment, amounting to 10.4±4.7 complexes on average (versus 59 complexes prior to HU). The high temporal resolution enabled us to determine the rate at which these complexes dissociate from forks, amounting to 15±6.9 complexes per min (Supplementary Table 3). Upon HU wash-out, PCNA reaccumulated to 33.7±16.6 complexes per fork. This incomplete re-accumulation can be attributed to cells exiting S shortly after HU wash-out. Applying the same approach to RPA1 quantification, we estimated that the number of RPA complexes engaged at a single fork peaked at 80.7±27.7 during the 1-h HU treatment period. Considering that the RPA1 heterotrimer occupies a region of approximately 30 nt [43–45], we calculated the total length of ssDNA generated following HU treatment, which peaked at 2421±832 nt per fork (Supplementary Table 2). Deriving the number of molecules engaged at forks at 30-s intervals allowed us to calculate the rates of ssDNA generation and RPA1 accumulation at 40±13.5 nt per min and 1.3±0.45 RPA complexes per min (Supplementary Table 3), respectively. HU wash-out resulted in an RPA removal rate of 8.9±5.2 complexes per min, which was approximately 7-fold faster than the rate of its accumulation.

### PCNA and RPA1 exhibit contrasting exchange rates at stalled forks

Once we quantified the number of PCNA and RPA complexes engaged at replication forks as well as their rates of recruitment and removal, we sought to determine the exchange rates of bound PCNA/RPA at stalled forks. To this end, we performed FRAP on individual PCNA/RPA1 foci 40 min after HU addition (Figure 4 and Figure S5). At this point, HU-induced PCNA removal is complete. We selected larger foci, in which there was a sufficient residual PCNA signal. FRAP of these foci revealed two phases of PCNA signal recovery (Figure 4D,E). The first was of rapid re-accumulation, reflecting freely diffusing molecules. The latter was more gradual, reflecting the exchange rate of replisome-bound PCNA. Fitting of the FRAP curve allowed us to clearly distinguish between these two fractions and specifically follow the recovery of fork-bound PCNA (Figure 4E,G,I). The half-time of PCNA signal recovery was 149±28.35 s, with approximately 30% of the bound fraction (∼3±1.3 complexes) being non-mobile (Supplementary Table 4, Figure 4I). Considering the number of residual complexes in HU-dissociated foci, we estimated that 2.4±0.9 PCNA complexes exchanged during the first minute after photobleaching (Figure 4J,K). For comparison, PCNA foci during normal replication (without HU treatment) (Figure 4C) exhibited much faster single-step complete recovery, with a half-time of 55±6.25 s (Figure 4I) and an exchange rate of 31.2±4.9 PCNA complexes per min. Our result demonstrates that exchange of PCNA at the stalled replication fork is significantly delayed and a considerable fraction remains non-mobile. This suggests that residual PCNA complexes persist on the leading strand and at least on some Okazaki fragments following fork stalling. As opposed to PCNA, RPA1 foci exhibited rapid single-step recovery at stalled replication forks, with a half-time of 5±0.95 s (Supplementary Table 4, Figure 4A,L). Within 9 s of photobleaching, 61±14 RPA complexes (out of ∼81 bound RPA complexes) exchanged, corresponding to a rate of 407.5±94 per min (Supplementary Table 4, Figure 4M), which indicates that, in contrast to PCNA, ssDNA-bound RPA at stalled replication forks is highly dynamic.

**Figure 4.**
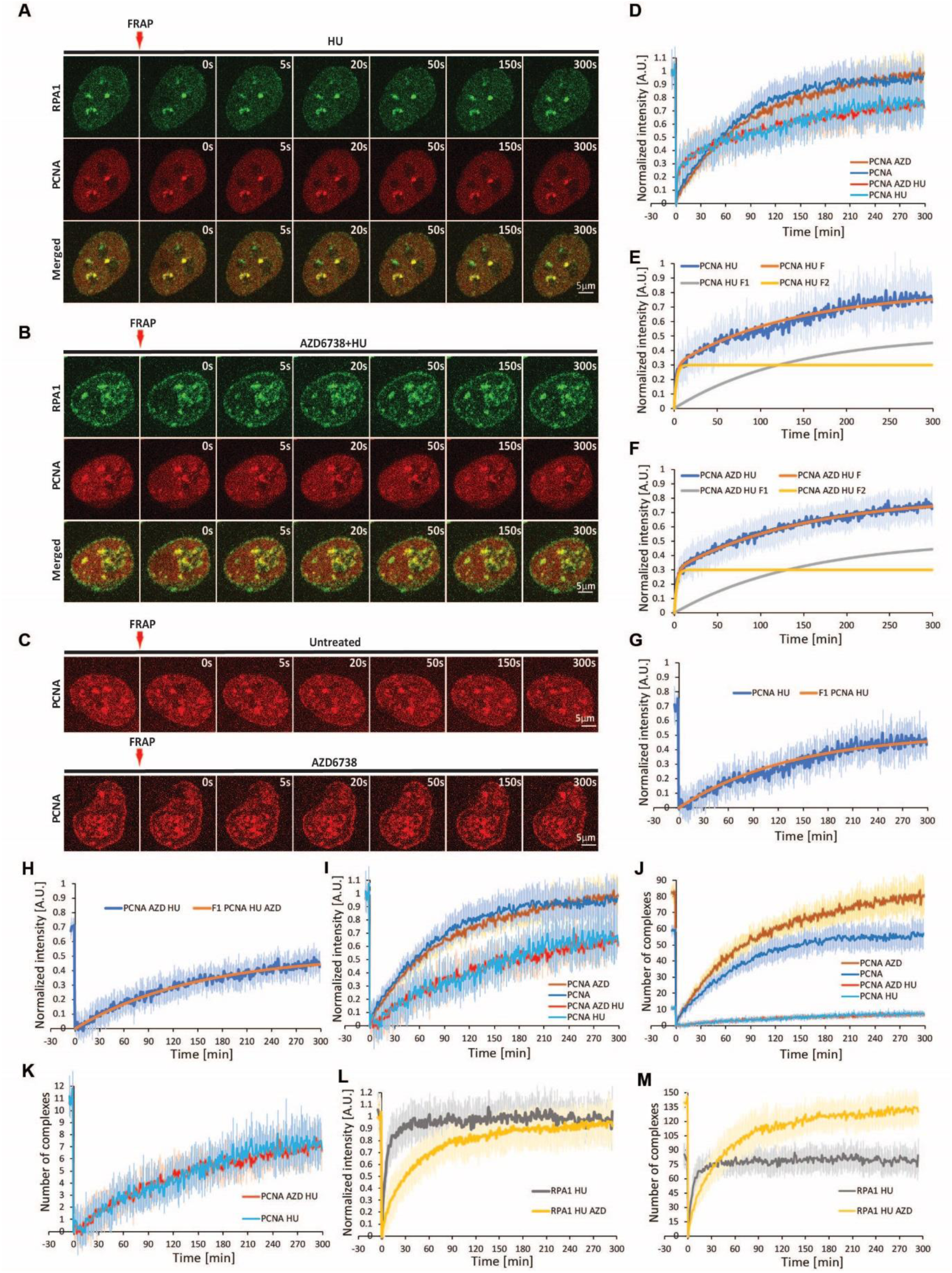
Exchange of PCNA and RPA1 at replication foci during unperturbed and stalled replication. (**a**) Representative timelapse images depicting simultaneous FRAP of RPA1 and residual PCNA co-localized at stalled replication forks under nucleotide depletion. (**b**) Same as (a), but under conditions of ATR inhibition. (**c**) Replication timelapse images of PCNA FRAP at replication foci during unperturbed replication (upper panel) or in the presence of ATR inhibitor AZD6738. (**d**) FRAP curves of PCNA under the following conditions: untreated, HU alone, AZD alone, HU+AZD. Mean intensity is normalized as described in Figure S5. (**e**) Contribution of distinct PCNA fractions (freely diffusing [F1] and replisome-bound [F2]) to the FRAP curve under nucleotide depletion, as determined via fitting of two single exponential curves. (**f**) Same as (e), but in the presence of both AZD6738 and HU. (**g**) Recovery of the replisome-bound fraction of PCNA under HU treatment, as derived based on (e). (**h**) Recovery of the replisome-bound fraction of PCNA under HU treatment, as derived based on (f). Comparison of PCNA recovery under the following conditions: untreated, AZD alone, HU alone, HU+AZD. The contribution of freely diffusing PCNA has been subtracted from the HU and HU+AZD curve, as per (g) and (h). (**j**) Number of PCNA complexes at a single replication fork, recovered after photobleaching. (**k**) Enlarged view of HU and HU+AZD curves from (j). (**l**) FRAP curves of RPA1 at replication foci under HU alone and HU+AZD. (**m**) Number of RPA complexes at a single replication fork recovered after photobleaching under HU alone and HU+AZD. For HU: n = 11 cells; for HU+AZD: n = 16 cells; for AZD (PCNA only): n =13 cells; untreated (PCNA only): n = 15 cells. **Abbreviations:** HU: hydroxyurea; AZD: AZD6738.

### ATR inhibition greatly enhances the amount and rate of RPA accumulation at stalled forks

Under conditions of replication stress, RPA coats the excess ssDNA generated at forks, facilitating the recruitment and activation of the ATR kinase, which acts to preserve, remodel, and restart stressed forks through the phosphorylation of replisome components and repair factors, while suppressing further origin firing [7,46,47]. To gain insight into the contribution of ATR to fork stalling and restart, we studied PCNA and RPA1 dynamics during HU treatment and wash-out under continued ATR inhibition (Figure 2B, Video 7). While PCNA dynamics were largely unaffected, a striking difference was observed in RPA1 recruitment, with a near-two-fold increase in RPA levels at replication foci (Figure 2C,D and Supplementary Table 1), accumulating with a 4-fold faster half-time (t_1/2_ = 5.6±2.2 min). This is in line with the well-established role of ATR in limiting helicase-polymerase uncoupling and preventing subsequent RPA exhaustion [15–18]. Importantly, RPA1 accumulation reached a plateau at approx. 25 min after HU treatment, engaging 70% of nuclear RPA1. Thus, beyond this timepoint, the concentration of freely diffusing RPA1 may no longer be sufficient to cover newly generated ssDNA. Employing the above-described conversions, we calculated that the total number of RPA complexes per fork increased to 139±13.7 (from 80.7), bound to approximately 4180±413 nt of ssDNA (Supplementary Table 2, Figure 2E,F). These equate to rates of 9.6±3.6 RPA complexes recruited per min and approximately 288±107 nt of ssDNA generated per min (Supplementary Table 3). Thus, ATR inhibition resulted in a 7.3-fold faster rate of HU-induced RPA1 loading. The total ssDNA generated under these conditions may considerably exceed this number since the RPA pool is exhausted as unwinding proceeds.

Following HU washout in the presence of ATRi, PCNA re-accumulation and RPA1 removal occurred at comparable rates to those observed without ATRi. However, RPA1 was not completely removed during fork restart, with approximately 16% remaining at replication foci (Figure 2D). This residual fraction persisted even after PCNA re-accumulation. An average of 22±16 RPA complexes and 660 nt of ssDNA per replication fork persist for 1 h after restart (Figure 2E,F).

FRAP analysis revealed a comparable exchange rate of the residual PCNA at HU-stalled forks, with versus without ATR inhibition (Figure 4I, K). Meanwhile, the RPA1 exchange rate was decreased six-fold, amounting to 34.2±9.8 complexes exchanged for the first 9 s after photobleaching, or a rate of 228.1±65.7 complexes/min (Supplementary Table 4, Figure 4L, M). Slow RPA1 exchange could reflect the exhaustion of the nuclear RPA1 pool and, possibly, the continuous generation of naked ssDNA, which is not coated by RPA1.

To gain further insight into cell fate following acute replication stress in the absence of ATR activity, we followed cells for 20 h (Video 7). As demonstrated above, cells subjected to HU-induced replication stress exhibited RPA1 accumulation at replication foci during S phase, suggesting ssDNA accumulation. However, this accumulation was resolved by G2, and cells proceeded through two mitoses, without post-replicative ssDNA (Figure S4A). In contrast, ATR inhibition combined with 1-h HU treatment resulted in the persistence of RPA-coated ssDNA into G2. Thereafter, the majority of cells died during mitosis or entered G1 with severely fragmented nuclei (Figure S4B, D). Taken together, our results demonstrate and quantify the drastic increase in the rate of RPA1 loading during replication fork stalling under ATR inhibition. Further, we show that ATR prevents post-replicative ssDNA gap persistence following fork restart and subsequent mitotic catastrophe.

### Co-inhibition of ATM and ATR prevents RPA removal during fork restart

Next, we evaluated the contribution of apical DDR kinase ATM to fork regulation under conditions of replication stress. To this end, cells were incubated with ATM inhibitor Ku55933 [48] prior to, during, and after HU treatment (Video 8). ATM inhibition did not have an effect on the dynamics nor on the amounts of PCNA and RPA1 during fork stalling and restart (Figure S6).Co-inhibition of both kinases, however, resulted in a slightly faster HU-induced RPA1 accumulation when compared to ATR inhibition alone (Figure 5D, Supplementary Table 3, Video 9). While the half-times and maximum levels of RPA1 recruitment in these two conditions were similar, RPA1 reached a plateau slightly faster under ATR and ATM co-inhibition. A notable difference was observed in the residual RPA1 following fork restart, with co-inhibition resulting in approximately 52% of RPA being retained versus 16% under ATR inhibition alone (Figure 5E, Supplementary Table 2). This amounted to approximately 74±18 RPA complexes covering ∼2200±540 nt of ssDNA (Figure 5H, I).

**Figure 5.**
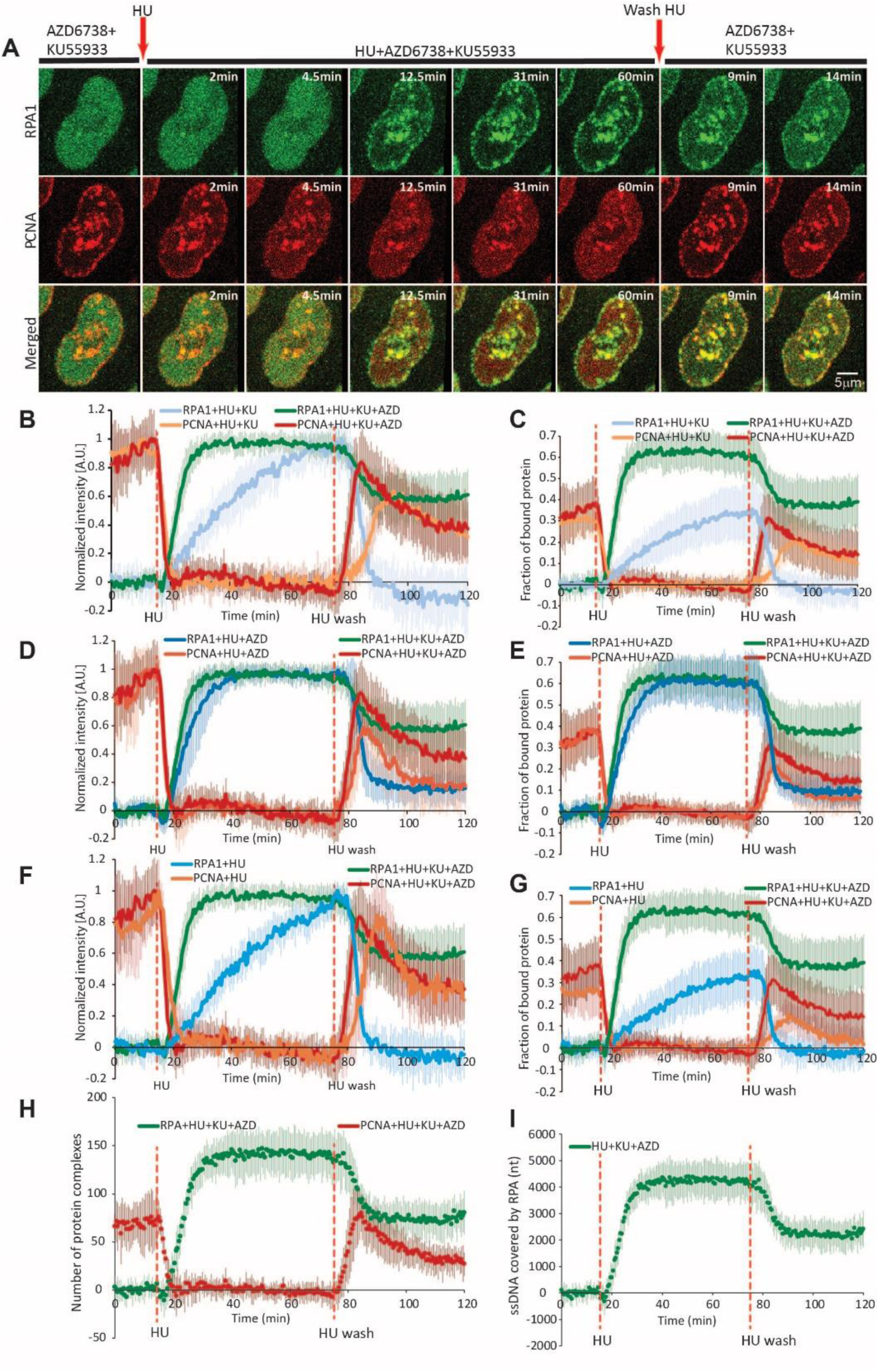
Influence of ATM activity on RPA1 and PCNA dynamics during HU-induced replication fork stalling and restart under conditions of ATR inhibition. (**a**) Representative time-lapse images of RPA1 and PCNA before, during, and after 10mM HU treatment under combined ATM (10µM Ku55933) and ATR (3µM AZD6738) inhibition. Arrows indicate timepoints of HU addition and washout. Scale bar = 5 µm. (**b**) Normalized kinetics of PCNA and RPA1 at replication foci during HU-induced replication fork stalling and restart under ATM inhibition with or without ATR co-inhibition. The maximum intensity of PCNA/RPA1 engaged at replication foci is normalized to 1. (**c**) Fraction of PCNA and RPA1 bound at replication foci during HU-induced replication fork stalling and restart under ATM inhibition with or without ATR inhibition, relative to the total nuclear intensity of PCNA/RPA1, which is normalized to 1. (**d**) Normalized kinetics of PCNA and RPA1 at replication foci during HU-induced replication fork stalling and restart under ATR inhibition with or without ATM co-inhibition. The maximum intensity of PCNA/RPA1 engaged at repl ication foci is normalized to 1. (**e**) Fraction of PCNA and RPA1 bound at replication factories during HU-induced replication fork stalling and restart under ATR inhibition with or without ATM inhibition, relative to the total nuclear intensity of PCNA/RPA1, which is normalized to 1. (**f**) Same as (b) and (d), but with or without combined ATR+ATM inhibition. (**g**) Same as (c) and (e), but with or without combined ATR+ATM inhibition. (**h**) Estimated number of PCNA homotrimer and RPA heterotrimer complexes engaged at replication foci during HU-induced fork stalling and subsequent restart. (**i**) Estimated number of nucleotides covered by RPA heterotrimers under combined ATM and ATR inhibition. Dashed orange lines indicate the timepoints of HU addition and wash-out. For HU+AZD+KU, n = 10 cells; for HU+KU: n = 19 cells. **Abbreviations**: HU: hydroxyurea; AZD: AZD6738; KU: Ku55933

To assess how persistent this large fraction of residual RPA is, we followed cells for 20 h under combined ATR and ATM inhibition (Video 10). Even after HU washout, the large residual RPA1 fraction persisted throughout S phase, through G2, and into mitosis, whereafter massive cell death occurred, with no cells reaching G1 (Figure S4C). Thus, when compared to ATR inhibition alone, ATM and ATR co-inhibition resulted in a considerable increase in HU-induced post-replicative ssDNA stretches and an inability to progress into the subsequent cell cycle. Taken together, our findings suggest that while ATM does not influence the rate of fork stalling in the presence of ATR activity, it prevents excessive post-replicative ssDNA accumulation in ATR-inhibited cells.

### PCNA and RPA dynamics are consistent among cell lines

We then assessed whether PCNA and RPA1 dynamics under conditions of HU-induced replication stress are consistent among cell lines. To this end, we generated PC3 and DU145 expressing mCherry-tagged PCNA and EGFP-tagged RPA1 via BAC recombineering, and subjected them to HU treatment alone or in combination with ATRi, ATMi, or both.

HU-induced PCNA removal occurred at identical rates among the three cell lines in all experimental conditions (Figure 6). Consistent with our results from HeLa, ATR inhibition considerably increased the rate of HU-induced RPA1 accumulation in PC3 and DU145 cells, while ATM inhibition had not significant effect on this process (Figure 6B,C). It should be noted that under all experimental conditions, RPA1 accumulation was slower in PC3 and DU145 than in HeLa. While RPA1 removal following HU washout occurred at comparable rates between the three cell lines, it began at later time points in the two former cell lines, especially in PC3, where a delay greater than 10 min relative to HeLa was observed (Figure 6A). Consistent with this, delays were also noted in PCNA re-accumulation after HU washout. In summary, while the overall dynamics of PCNA and RPA1 were consistent among the three cell lines, differences were observed in RPA1 accumulation and removal, particularly in PC3. This could stem from differences in nucleotide metabolism, for example, slower recovery of the nucleotide pool.

**Figure 6.**
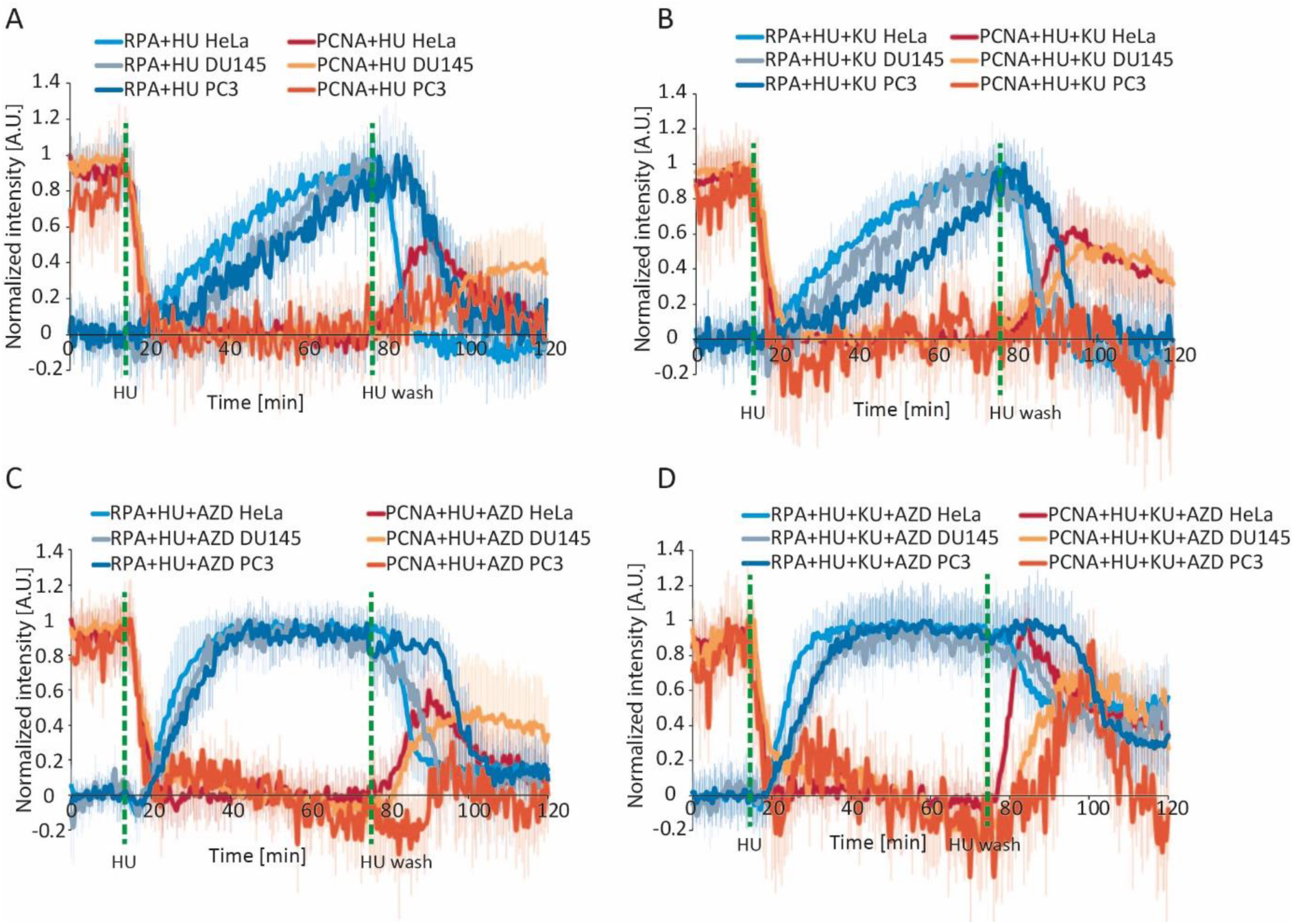
Dynamics of PCNA and RPA1 during hydroxyurea-induced replication stress in HeLa, DU145, and PC3 cell lines. (**a**) Normalized intensity of PCNA and RPA1 during, before, and after HU treatment. (**b**) Same as (a), but under conditions of ATM inhibition (10µM Ku55933). (**c**) Same as (a), but under conditions of ATR inhibition (3µM AZD6738). (**d**) Same as (a), but under combined ATR and ATM inhibition. Dashed green lines indicate the timepoints of HU addition and washout. Data are presented as the mean±SD. For HeLa: n = 11 cells (HU), n = 16 cells (HU+AZD), n = 19 cells (HU+KU), n = 10 cells (HU+AZD+KU); for DU145: n = 11 cells (HU), n = 10 cells (HU+AZD), n = 10 cells (HU+KU), n = 18 cells (HU+AZD+KU); for PC3: n = 18 cells (HU), n = 10 cells (HU+AZD), n = 13 cells (HU+KU), n = 15 cells (HU+AZD+KU). **Abbreviations**: HU: hydroxyurea; AZD: AZD6738; KU: Ku55933

### MRE11-mediated resection does not influence RPA dynamics during fork stalling and restart

As ssDNA at stressed forks can be generated via nucleolytic degradation, we sought to assess whether MRE11, which is known to mediate pathological nascent strand degradation at stalled forks, would affect RPA and PCNA dynamics upon nucleotide depletion. To this end, we inhibited MRE11 exonuclease activity with mirin [49] prior to HU-induced stalling and release (Video 11). Treatment with mirin alone had no effect on PCNA and RPA1 dynamics during fork stalling (Figure S7). Further, no changes were noted in PCNA-reaccumulation and RPA1 removal during fork restart. To assess whether MRE11 contributes to the increased speed and levels of HU-induced ssDNA accumulation during ATR inhibition, we subjected cells to combined mirin and ATRi treatment prior to HU (Video 12). Once again, no difference in PCNA/RPA1 dynamics were noted (Figure S8). These results are in line with the notion that forks are largely protected from MRE11-mediated nucleolytic degradation in BRCA1/2-competent cells[21].

### RAD18 dynamics at stalled forks are not influenced by ATR activity

The RAD18 E3 ubiquitin ligase is recruited to stalled forks where it interacts with and monoubiquitinates PCNA to facilitate damage tolerance [50]. Further, RAD18 is implicated in the repair of DSBs within replicated chromatin [51]. In light of this, we set out to measure the dynamics of RAD18 recruitment and removal upon HU-induced replication stress (Video 13). RAD18 foci were observed during unperturbed replication, but did not always co-localize with PCNA foci. Upon HU treatment, RAD18 was rapidly recruited to some replication sites, which coincided with the dissociation of these pre-existing RAD18 foci (Figure 7A). The half-time of RAD18 recruitment at replication foci was comparable to that of PCNA removal (t_1/2_= 4.5 min versus 2.5 min) and thus considerably faster than RPA accumulation (Figure 7B). RAD18 reached a plateau at approx. 14.5 min after HU treatment. In contrast to RPA1, RAD18 foci tended to persist even after PCNA re-accumulation during fork restart. However, our approach of measuring the changes in freely diffusing protein outside of foci was not suitable for characterizing RAD18 dynamics because previously existing dissolved, while new foci simultaneously formed at replication factories upon HU treatment. Tracking of individual PCNA and RAD18 foci via SPARTACUSS confirmed that, after HU addition, the latter was recruited at least in some replication foci, with a half-time comparable to that of PCNA removal (Figure 7C). This was observed with or without ATR inhibition (Figure 7D). Importantly, individual tracking confirmed that the above-described pre-existing RAD18 foci, dissolved at a comparable rate to that of PCNA foci. Our findings highlight that upon acute replication stress, RAD18 is rapidly redistributed from non-actively-replicating regions, which likely harbor DNA lesions, to sites of fork stalling, where it tends to persist even after restart. Finally, ATR inhibition had no effect on RAD18 recruitment and persistence at replication foci (Video 14).

**Figure 7.**
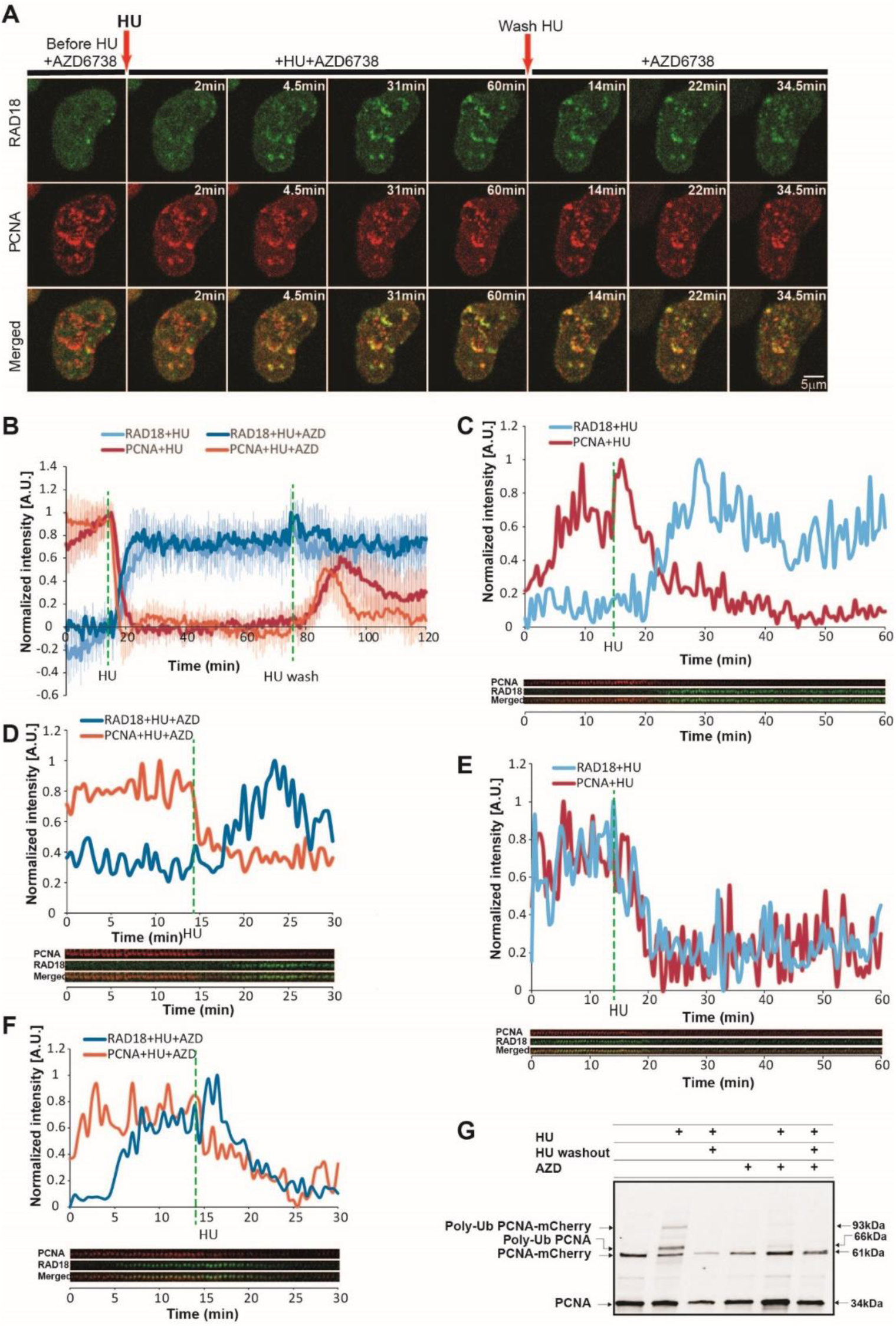
Dynamics of RAD18 during hydroxyurea-induced replication fork stalling and restart. (**a**) Representative time-lapse images of RAD18 and PCNA before, during, and after HU treatment. Arrows indicate timepoints of HU addition and washout. Scale bar = 5 µm. (**b**) Normalized kinetics of RAD18 and PCNA during HU-induced replication fork stalling and restart, with or without ATR inhibition (3µM AZD6738). The maximum intensity of PCNA/RAD18 foci is normalized to 1. (**c**) Normalized intensity of single RAD18 and PCNA foci before and after HU addition (max intensity = 1). Single foci were tracked using SPARTACUSS, and a representative kymogram is shown. This panel presents a case where RAD18 accumulates at a replication focus while PCNA dissociates after HU treatment. (**d**) Same as (c), but under conditions of ATR inhibition. (**e**) Same as (c), but this panel presents a case where RAD18 is already present at the replication focus and dissociates from the replication focus in parallel to PCNA upon HU addition. (**f**) Same scenario as shown in (e), but under conditions of ATR inhibition. Dashed green lines indicate timepoints of HU addition and removal. (**g**) Western blot analysis of PCNA under conditions of acute HU-induced replication stress, with or without ATR inhibition. The appearance of multiple PCNA bands reflected the generation of poly-ubiquitinated forms of both the endogenous and BAC-tagged PCNA. For HU: n = 17 cells; for HU+AZD: n = 12 cells. **Abbreviations**: HU: hydroxyurea; AZD: AZD6738

RAD18 monoubiquitylates PCNA to enable translesion synthesis in the face of replication fork barriers. This monoubiquitylation at K164 can then be extended via K63-linked polyubiquitylation, facilitating DNA damage tolerance through other mechanisms [52]. Thus, we assessed PCNA ubiquitination through immunoblotting. Our results demonstrated that HU treatment gave rise to a two distinct bands, which were ∼30kDa heavier than the endogenous and mCherry-fused PCNA, respectively (Figure 7G). This corresponded to PCNA with four ubiquitin molecules attached. The small amount of polyubiquitylated PCNA aligns with our observation that a minor fraction of PCNA molecules remain at replication foci during HU-induced fork stalling. The observed polyubiquitylation suggests that a tolerance mechanism other than monoubiquitylation-stimulated translesion synthesis is at play during nucleotide depletion. ATR inhibition considerably suppressed HU-induced PCNA polyubiquitylation, suggesting that ATR activity may be, at least in part, required for this modification to occur.

## Discussion

In the present work, we employed live-cell imaging to study the dynamics of PCNA and RPA1 during replication fork stalling and subsequent restart at an unprecedented temporal resolution in single cells. Our results provide a detailed timescale and rates for the major molecular processes during the replication stress response, namely, PCNA turnover and ssDNAgeneration (Figure 8).

**Figure 8.**
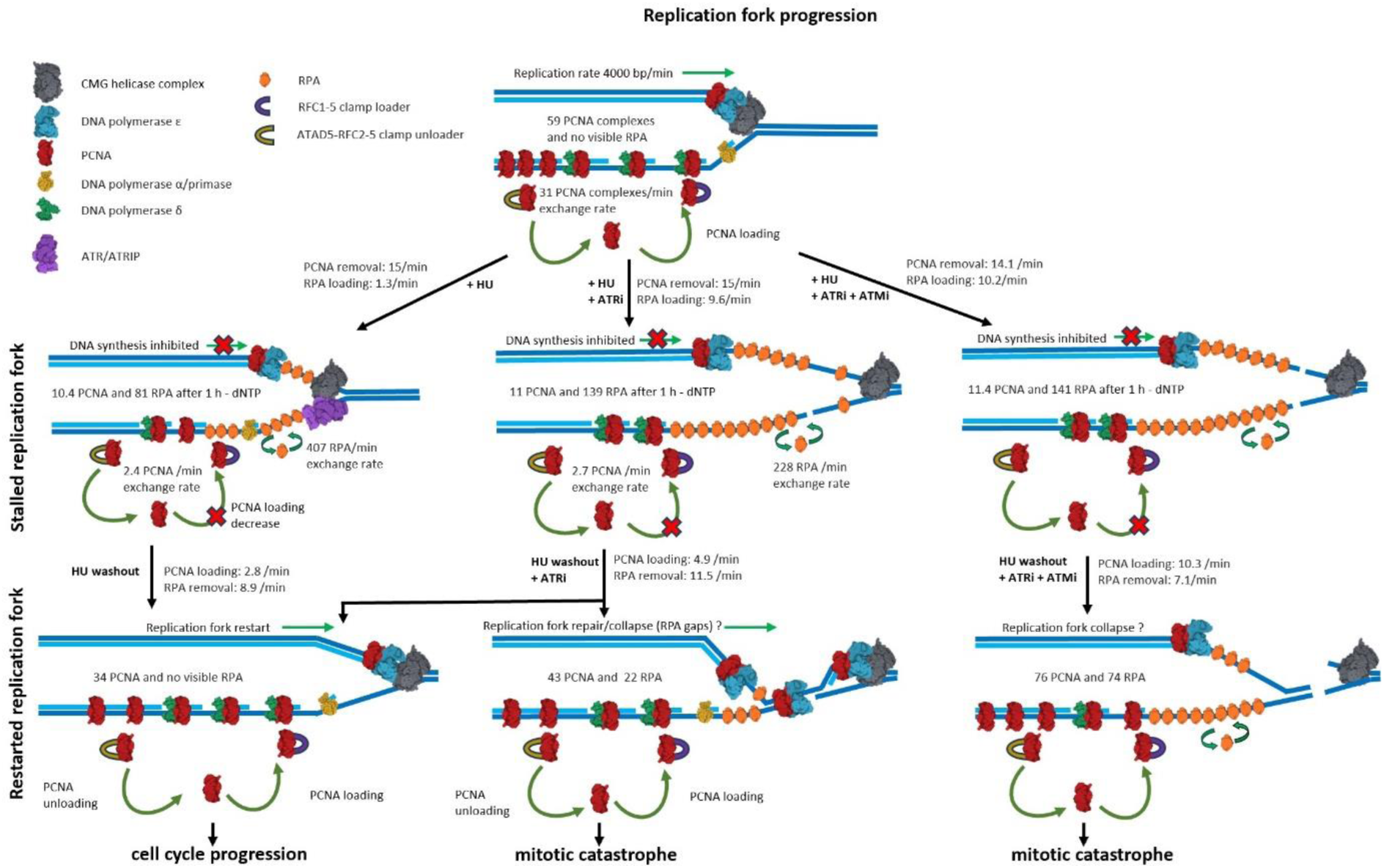
Visual summary of PCNA and RPA dynamics during replication fork stalling and restart.

Finally, we characterize how perturbing the intra-S checkpoint and key DDR kinase activity influences the above-described kinetic measures.

Our findings show that the majority (85%) of PCNA engaged at replication foci is rapidly removed following HU treatment. The amount of PCNA at the fork depends on its recruitment to the leading and lagging strands by the RFC clamp loader complex as well as its removal by ATAD5-RLC upon completion of DNA synthesis [32]. Thus, the removal of PCNA that we observe can be attributed to a decrease in PCNA loading due to nucleotide depletion and the continuous removal of previously engaged PCNA after the completion of lagging strand synthesis.

The kinetics of RPA during nucleotide depletion reveal that despite an active intra-S checkpoint, RPA loading gradually proceeds for up to 1.5 h. The prevailing view posits that the accumulation RPA-coated ssDNA triggers ATR activation, leading to the phosphorylation of replisome components, including Claspin, RPA, and MCM subunits, which limits DNA unwinding to prevent further ssDNA accumulation and RPA exhaustion [7,18]. Our measurements indicate that, while ATR slows the rate of ssDNA accumulation about nine-fold, the amount of RPA recruited at forks after 1 h increases two-fold. However, the outcomes drastically differ, as ssDNA persists into G2 in ATR-inhibited cells, which then undergo mitotic catastrophe, as opposed to the smooth cell cycle progression observed in non-inhibited cells. Inhibition of MRE11 exonuclease activity had no effect on RPA1 recruitment, which aligns with BRCA-dependent fork protection against pathological MRE11-mediated nascent strand degradation [21]. Therefore, we attribute the rapid and extensive RPA1 accumulation to uncoupling of replicative unwinding from DNA synthesis under ATR inhibition. More importantly, our results demonstrate that while the ATR-dependent intra-S checkpoint severely slows unwinding, it does not prevent considerable ssDNA accumulation over time. We sought to compare these rates to the rate of ssDNA generation that we determined based on RPA1 recruitment during nucleotide depletion. When ATR is inhibited, ssDNA was generated at a rate of 290 nt per min (4.8 nt per s). Considering that ssDNA probably accumulates on both strands, DNA is being unwound at a two-fold slower rate, namely, 2.4 bp per s. Intriguingly, this value is very close to the in vitro single-molecule measures of unwinding at 6.1 bp per s by the drosophila Cdc45/Mcm2–7/GINS (CMG) helicase [53,54] and 5-10 bp per s by the yeast CMG [55]. The similar rates of *in vitro* unwinding and *in vivo* ssDNA generation under ATR inhibition further support uncoupling as the driver of RPA1 recruitment.

While our findings do not exclude fork reversal, the continuous generation of ssDNA during 1-h fork stalling is hard to reconcile with reversed fork structures, which should, in theory, reduce ssDNA as a result of nascent strand reannealing.

Taking advantage of the temporal resolution provided by our system, we sought to assess the contribution of ATM to replication fork stalling and reversal upon nucleotide depletion. While ATM inhibition had no effect on PCNA and RPA dynamics during fork stalling and restart under ATR proficiency, co-inhibition of both revealed that ATM largely prevents the persistence of ssDNA after S-phase when ATR is inhibited. The exact mechanism through which ATM suppresses these ssDNA stretches is currently unclear. One plausible explanation would be that the rapid and extensive unwinding that occurs during ATR inhibition leads to the generation of ssDNA stretches that are not protected by RPA and are thus susceptible to DSB generation. ATM-driven repair would contribute to the resolution of these lesions, reducing post-replicative RPA.

Interestingly, the rapid PCNA removal coincided with RAD18 recruitment within the same replicating regions. This was rather unexpected considering the well-established role of RAD18 in PCNA monoubiquitylation during translesion synthesis [56]. One possible explanation may be that RAD18 monoubiquitylates the remaining fraction (16%) of PCNA present within replication foci. In principle, nucleotide depletion should not elicit translesion synthesis nor template switching, both of which require the presence of unrepaired DNA lesions. While RAD18 is implicated in DSB repair, it is unlikely that DSBs accumulate as early as 5-10 min after HU addition, especially in the context of active ATR signaling. It was previously demonstrated that RAD18 binds RPA at the replication fork [50], which could explain the RAD18 recruitment dynamics that we observe at the stalled replication forks. However, it should be noted that HU-induced RAD18 accumulation occurred at a much faster rate than that of RPA, and the 10-fold increase in RPA1 loading under ATR inhibition did not affect RAD18 dynamics. These results do not exclude an RPA-dependent mechanism for RAD18 loading at stalled forks, but rather suggest that a small amount of initially loaded RPA may be sufficient.

Our results demonstrated that ATR inhibition suppressed HU-induced PCNA polyubiquitylation, which is somewhat counterintuitive considering that RAD18 accumulated in identical fashion with or without ATR inhibition. Furthermore, RAD18 foci persisted after fork restart, regardless of ATR activity. We also did not detect PCNA monoubiquitylation, the major function ascribed to RAD18. These results can be explained by a model whereby monoubiquitylated PCNA, which is unable to proceed along the template due to nucleotide depletion, rather than DNA lesions, is rapidly polyubiquitylated [52].

In summary, the current work provides a highly detailed view into the dynamics of replication fork stalling and restart during nucleotide depletion. The live-cell imaging-based approach proposed herein, combined with advanced image analysis, enables us to clearly distinguish two phases of fork stalling. First is the dissociation of most PCNA molecules from the replisome, which reflects the abrupt drop in DNA synthesis. This is followed by a gradual increase of RPA-coated ssDNA stretches despite an active intra-S checkpoint and regardless of MRE11 exonuclease activity. ATR inhibition drastically increases the rates of ssDNA generation, which aligns with previous in vitro measurements of DNA unwinding by eukaryotic replicative helicases. Crucially, our method enabled us to follow fork stalling and subsequent restart at the same replication factories within individual cells. Quantifying the rate of HU-induced RPA1 removal during fork restart revealed no differences with versus without the inhibition of ATR, ATM, or both. Our results indicate that ATR, but not ATM, is required for the resolution of ssDNA generated during hydroxyurea-induced replication stress. However, co-inhibition revealed that, in the absence of ATR activity, ATM prevents the persistence of vast ssDNA stretches upon fork restart. The approach developed herein enables one to follow major processes during replication fork stalling and restart with unprecedented temporal resolution, providing a basis for studying the effects of anticancer agents, which target replication stress.

## Materials and methods

### Cell culture

HeLa Kyoto, PC-3, and DU145 cells were used in the current work. HeLa Kyoto cells were cultured in Dulbecco’s Modified Eagle Medium (DMEM) supplemented with 10% fetal bovine serum (FBS), 100 units/ml penicillin, and 100 mg/mL streptomycin at 37°C and 5% CO2. PC-3 and DU145 were cultured in RPMI 1640 medium containing 2mM L-glutamine and 25mM sodium bicarbonate, supplemented with 10% FBS, 100 units/ml penicillin, and 100 mg/mL streptomycin. Cells at approx. 20% confluence were seeded into 35-mm MatTek glass-bottom dishes at least 48 h prior to imaging. For live-cell imaging of HeLa Kyoto cells, the culture medium was replaced with FluoroBrite DMEM (Thermo Fisher Scientific) supplemented with 10% FBS, 100 units/ml penicillin, 100 mg/mL streptomycin, and 2mM GlutaMAX. PC-3 and DU145 cells were imaged in their culture medium.

### Image acquisition and treatment

#### Treatment

Cells were treated with inhibitors of ATM (10µM KU55993), ATR (3µM AZD6738), MRE11 (100µM Mirin), alone or in combination, prior to imaging. Multiple fields were selected based on the presence of currently replicating cells, i.e. cells with discernible PCNA foci. Cells were imaged for 15 min prior to the addition of HU to the medium. All inhibitors were dissolved in DMSO, with the exception of HU, which was dissolved directly in the medium. The final concentration of DMSO in medium did not exceed 0.1%. The FluoroBrite DMEM (Thermo Fisher Scientific) media used for all treatment and washout steps were pre-heated to 37°C.

#### Timelapse spinning-disk confocal imaging

Images were acquired on a Andor Dragonfly 500 (Oxford Instruments, Oxfordshire, UK), equipped with a Nikon Eclipse Ti-E (Nikon, Tokyo, Japan) inverted microscope, a Nikon Perfect Focus System (PFS) (Nikon, Tokyo, Japan), a Nikon CFI Plan Apo 60x (NA 1.4) (Nikon, Tokyo, Japan) water immersion objective, and an iXon888 EMCCD (Oxford Instruments, Oxfordshire, UK) camera. Five or six fields were selected based on the presence of S-phase cells and followed in parallel. The cells were imaged in three focal planes with a step size of 0.5 μm, at intervals of 30 sec. For every experimental condition, the following timeframes were followed: (a) cells were imaged for 15 min; (b) HU was added, and cells were imaged for another 1-3 h movie; (c) HU was washed out, cells were imaged for 2 h thereafter. Whenever ATM, ATR, or MRE11 inhibitors were used, these were added 1 h prior to HU treatment and were constantly present in the medium (also included in the pre-warmed medium used to wash cells from HU).

For overnight timelapse imaging, cells were treated as described above and imaged every 10 min. With regard to acquisition, cells treated with HU alone were imaged once prior to HU addition, then followed for 1 h under HU treatment, and, finally, followed over approx. 20 h.

#### Airyscan imaging

For airyscan imaging, we used a Zeiss LSM980 with Airyscan 2 equipped with a 63x Plan-Apochromat (1.4NA) oil immersion objective. Cells were imaged every 2 min and were subjected to HU treatment as described above, but without wash-out. AZD6738 was added 1 h prior to imaging and was present in the media used for HU treatment. The cells were imaged in seven focal planes with a step size of 0.17 μm.

#### FRAP

To determine the exchange rates of PCNA and RPA1 at replication foci, we performed FRAP on an Andor Revolution spinning disk confocal system with a Nikon Eclipse Ti-E inverted microscope equipped with the Nikon Perfect Focus System (PFS), a Nikon CFI Plan Apo VC 60x (NA 1.2) water immersion objective, an iXon897 EMCCD camera, and an Andor FRAPPA module. We imaged cells in both the 488 and 561 nm channels every 1 s, including 5 frames prior to photobleaching, followed by 300 s. Photobleaching of both EGFP-RPA1 and mCherry-PCNA was performed using the 488nm laser with a nominal power of 30mW attenuated to 30% with 20 repeats and 100ms dwell time.

### Image analysis

All time-lapse imaging data were analyzed using the CellTool software [57]. Prior to analysis, a maximum intensity projection (of the three z-slices) was generated. All cell nuclei were segmented, selected as regions of interest (ROI), and tracked. Each cell was then saved as a separate file and subjected to a second registration in order to minimize cell movement within the cropped film. This secondary registration was performed using the MultiStackReg in ImageJ [58]. The proteins studied in the current work form foci within the nucleus in accordance with their function (e.g. PCNA accumulates at replication factories, RPA accumulates at ssDNA). These foci are often very abundant, close to each other, and dynamic, which together complicates their segmentation and simultaneous tracking. To measure the accumulation and removal of proteins at nuclear foci, we selected three ROIs in nuclear regions devoid of foci. The intensity of these foci-free ROIs corresponds to the amount of free protein within the nucleus and thus increases or decreases in this amount of free protein due to its recruitment and removal to chromatin reflect the changes in the chromatin-associated fraction. We ensured that these ROIs did not contain foci at any point during the time-lapse. A background ROI outside of the nucleus was also selected. The intensity measured at this ROI was subtracted from the intensities of all other ROIs in order to remove the background. See the formulas in Figure 3 for a detailed overview of the mathematical procedures used for the quantification and normalization performed in the current work. These formulas were based on several constants, including the number of RPA1 molecules/RPA complexes in the cell, the number of PCNA homotrimers in the cell, an average number of active replication forks at a given timepoint during S phase [29], and the number of nucleotides bound by a single RPA complex [43–45].

#### Analysis of FRAP experiments

Regions of interest included the cell nucleus, a replication focus that is photobleached, a replication focus that is not photobleached, and a region outside of cells. Constants and formulas for the calculation of variables used for quantifying the kinetics of RPA1 and PCNA exchange are presented in Figure S4. In brief, we: (a) subtract the noise from each of the two replication foci; (b) normalize the mean intensity of the six frames before photobleaching to 1 (for both foci); (c) divide normalized intensity of the photobleached focus by that of the non-photobleached focus; (d) we normalize the intensity value immediately after photobleaching to 0, and, finally, we once again normalize the average of the six frames before photobleaching to 1. From the resulting FRAP curve, we derive the half-times of recovery.

To determine the rates of recovery in molecules/complexes rather than mean intensities and half-times, we multiply the normalized intensity obtained in step (d) by the number of molecules at a single fork right before photobleaching, which we estimated for each experimental condition. To obtain the molecule exchange rate, we take a 9-s and 1-min interval from the FRAP curves for RPA1 without and with AZD6738, respectively. These represent the steepest parts of the recovery slope.

#### Analysis of single foci

Individual foci were tracked in Fiji using the MTrackJ plugin (ImageScience). The exported coordinates were used for further processing in SPARTACUSS, a software tool we previously developed [42]. Specifically, for each time point, we crop a 7×7 square around the focus center and the resulting windows are stitched together in what we refer to as a kymogram. The mean fluorescence inside a circle of radius 3.5 pixels centered at the focus is recorded for each time point.

### Western blot

Cells co-expressing RPA1-EGFP and PCNA-mCherry were seeded into six-well plates and cultured in DMEM containing 10% FBS, 400 µg/mL geneticin, and 2 µg/mL blasticidin. Cells were treated with the following combinations of inhibitors: HU, HU plus AZD6738, HU plus KU55933, AZD6738 plus KU55933, or DMSO. The final concentrations of HU, AZD6738, and KU55933 were as 10mM, 3µM, and 10µM. Cells were incubated with inhibitor for 90 min, washed with PBS, and lysed in RIPA lysis buffer (0.9% NaCl, 55mM Tric-HCl, 0,1% Triton, 0.5% Na2C2O4) containing a protease inhibitor cocktail. After a 20-min incubation at 4°C, cells were homogenized using an ultrasonic homogenizer. The resulting lysates were centrifuged for 20 min at 1200 rpm at 4°C. The obtained total protein was separated via 8-16% SDS-PAGE and transferred onto nitrocellulose membranes. After blocking, membranes were incubated with mouse primary antibodies against PCNA (PC10) (sc-56, Santa Cruz Biotechnology) and RPA70/RPA1 (MBS600132, MyBioSource) overnight at 4°С, followed by incubation with an IRDye® 800CW Goat anti-Mouse IgG Secondary Antibody (926-32210, LICORbio) at room temperature for 1 h. Bands were visualized using an Odyssey Infrared Imaging System and saved as TIF images. Band intensities were determined and compared using ImageJ.

## Acknowledgements

We would like to thank Sylvia Marinova-Varhoshkova and Joana Kicheva for critically reading the manuscript as well as Petar Takov for assistance with data analysis.

## Author contributions

Conceptualization, S.S.S.; methodology, S.S.S.; software, G.D.; formal analysis, T.D.-D., G.D., P.-B.V.K.; investigation, T.D., S.U., P.-B.V.K., A.A., A.I.; data curation, S.S.; writing— original draft preparation, P.-B.V.K., S.S.S.; writing—review and editing, P.-B.V.K., S.S.S.; visualization, T.D., G.D., A.A., R.S., P.-B.V.K., S.S.S.; supervision, S.S.S.; project administration, S.S.S.; funding acquisition, S.S.S. All authors have read and agreed to the published version of the manuscript.

