## Supplementary_Materials for "In and out of replication stress: PCNA/RPA1-based dynamics of fork stalling and restart in the same cell": Supplementary_Materials.pdf

### Supplementary Information

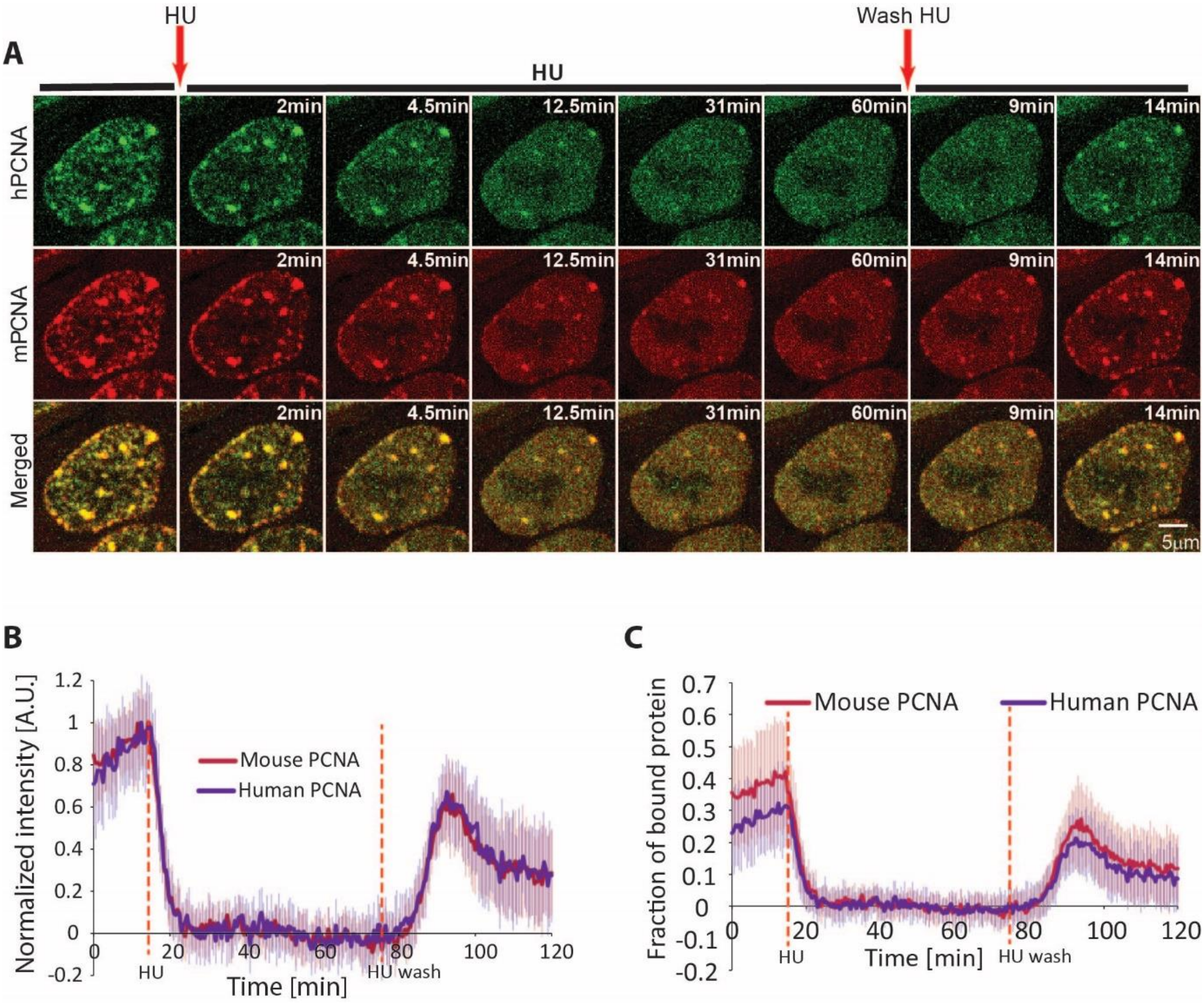


**Figure S1. Comparison of mouse versus human PCNA dynamics in response to hydroxyurea-induced replication stress.** (**a**) Representative time-lapse images of human PCNA (hPCNA-EGFP) and mouse PCNA (mPCNA-mCherry) before, during, and after 10mM HU treatment. Arrows indicate timepoints of HU addition and washout. (**b**) Normalized kinetics of mouse and human PCNA at replication foci during HU-induced replication fork stalling and restart. The maximum intensity of PCNA/RPA1 engaged at replication foci is normalized to 1. (**c**) Fraction of mouse and human PCNA bound at replication foci during HU-induced replication fork stalling and restart. Dashed orange lines indicate the timepoints of HU addition and wash-out. N = 21 cells. **Abbreviations**: HU: hydroxyurea

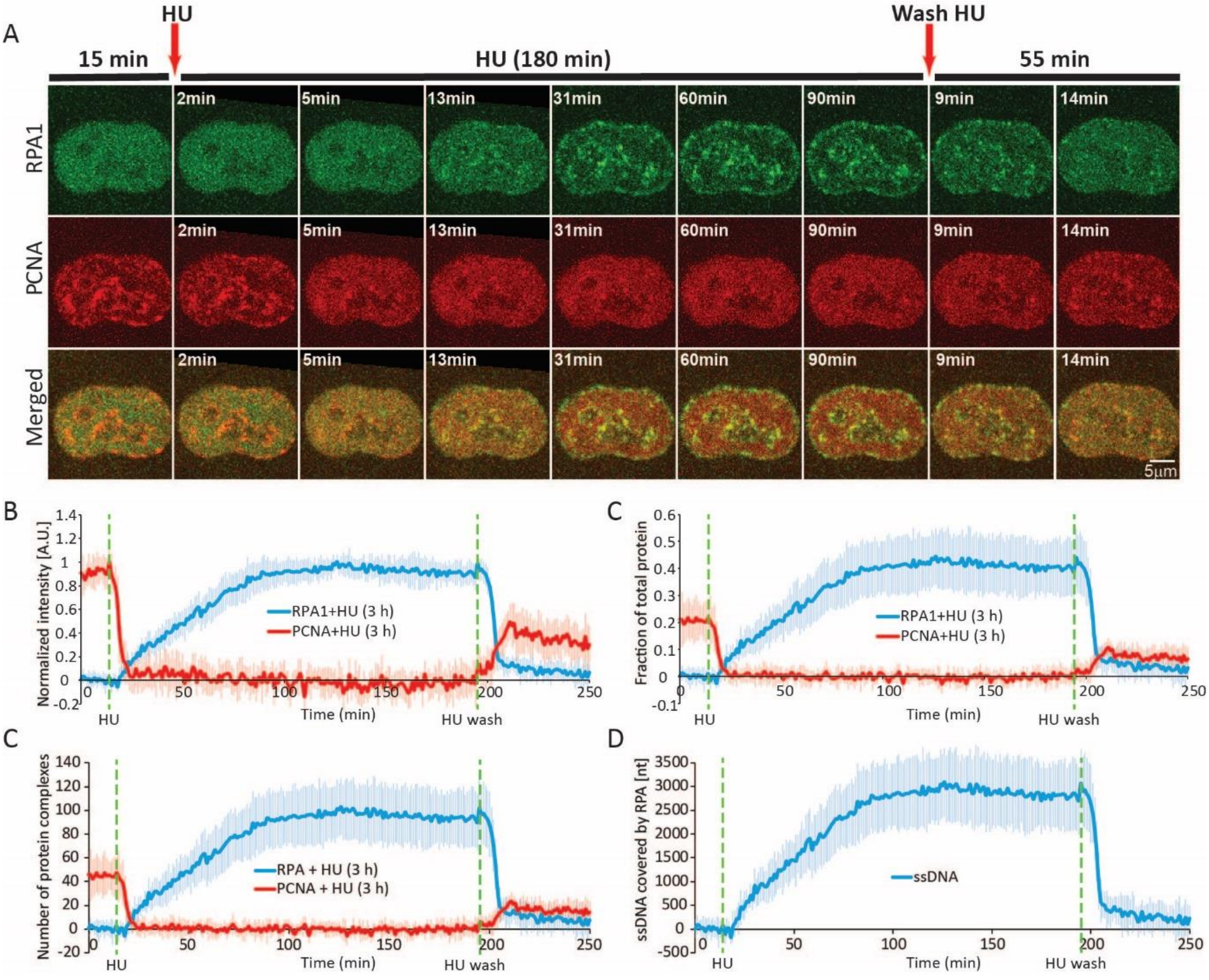


**Figure S2. PCNA and RPA1 dynamics during prolonged fork stalling under HU-induced nucleotide depletion.** (**a**) Representative timelapse images of RPA1-EGFP and mCherry-PCNA before, during, and after 3-h HU treatment. Timepoints of HU (10mM) addition and washout are indicated by red arrows. (**b**) Normalized kinetics of PCNA and RPA1 at replication foci during HU-induced replication fork stalling and restart. (**c**) Fraction of PCNA and RPA1 bound at replication foci during HU-induced replication fork stalling and restart, relative to the total nuclear intensity of PCNA/RPA1, which is normalized to 1. (**d**) Average number of RPA and PCNA complexes engaged at a single replication fork. (**h**) Average length of ssDNA (nt) covered by RPA at a single replication fork. Dashed green lines indicate the timepoints of HU addition and washout. Data are presented as mean ± SD. N = 11 cells. **Abbreviations**: HU: hydroxyurea

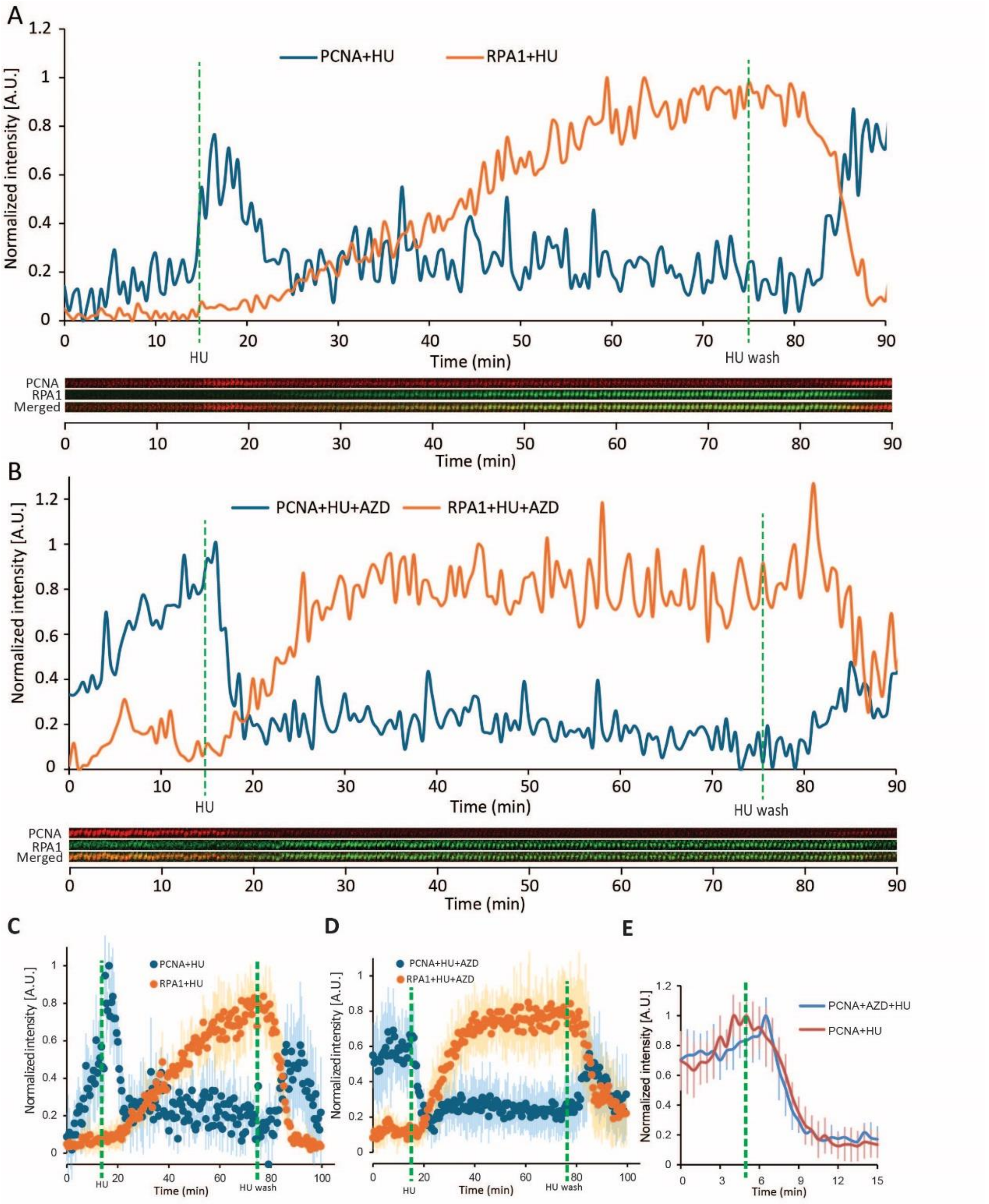


**Figure S3. Single focus dynamics during hydroxyurea-induced fork stalling and restart.** (**a**) Normalized kinetics of single PCNA and RPA1 foci during HU-induced replication fork stalling and restart. The maximum intensity of foci is normalized to 1. Single foci were tracked using the SPARTACUSS software tool (see Materials and methods), visualized via a kymogram (below). (**b**) Same as (a), but in conditions of ATR inhibition (3µM AZD6738). (**c**) Normalized intensity of single PCNA and RPA1 foci before, during, and after HU treatment. (**d**) Same as (**c**), but under ATR inhibition. Data are presented as mean±SD. (**e**) Dynamics of single PCNA foci dissolution upon HU addition, with and without ATR inhibition. Data are presented as mean±SD. Dashed green lines indicate timepoints of HU addition and wash-out. For HU: n = 4 foci; for HU+AZD = 14 foci. **Abbreviations**: HU: hydroxyurea; AZD: AZD6738

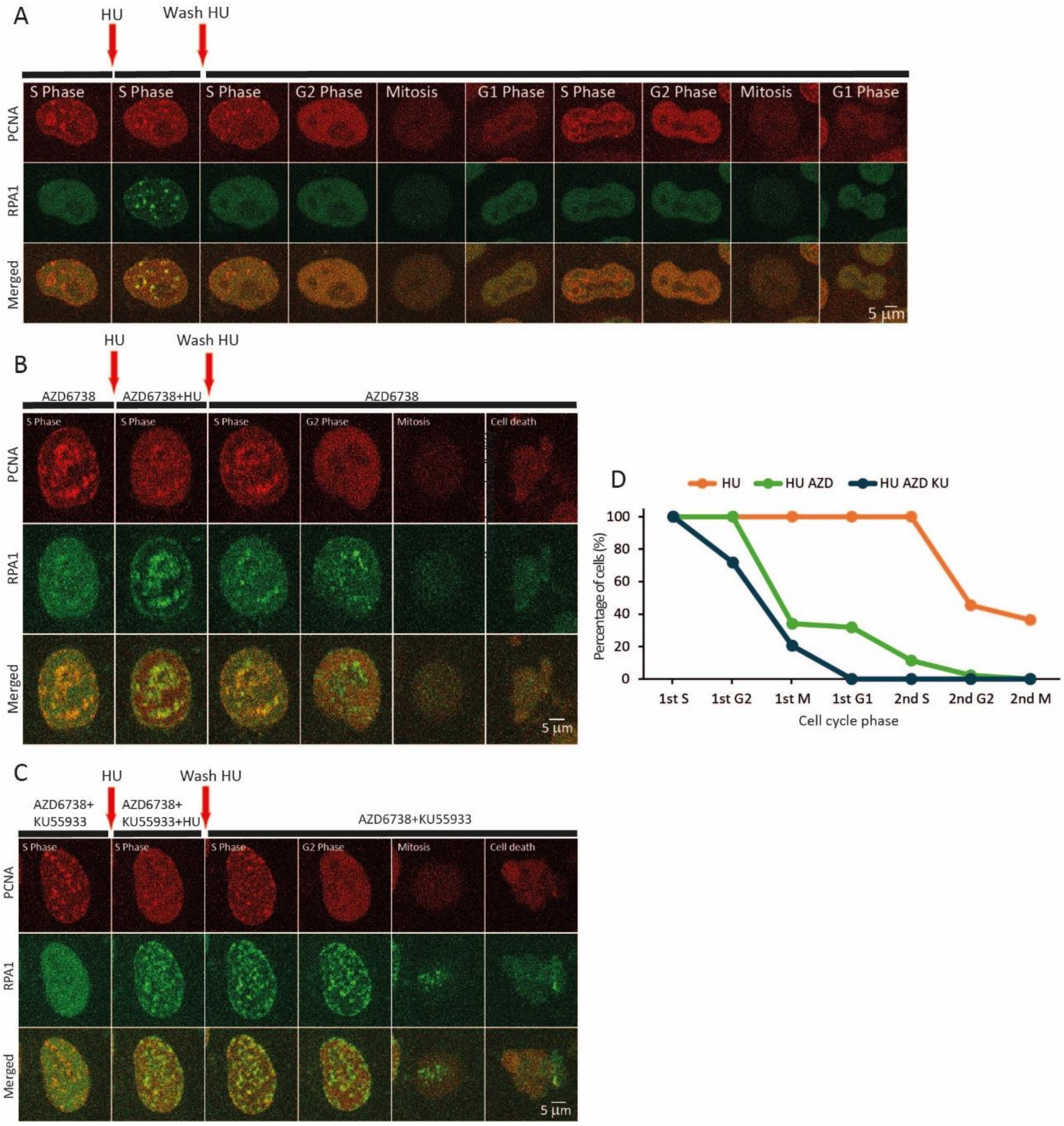


**Figure S4. Methodology for FRAP analysis of PCNA and RPA1 at replication foci.** (**a**) Regions of interest applied for analysis. 'A' represents the cell nucleus (white lining), 'B' represents a replication focus that is photobleached (blue ellipse), 'C' represents a replication focus that is not photobleached, used for normalization (orange ellipse), and 'D' represents the noise outside of cells (yellow ellipse). (**b**) Constants and formulas for the calculation of variables used for quantifying the kinetics of RPA1 and PCNA exchange. (**c**) Mean intensity of RPA1-EGFP within regions 'B' and 'C', calculated by subtracting constant $D_t$ from $B_t$ and $C_t$, respectively. (**d**) Normalized intensity of RPA1-EGFP/mPCNA-mCherry bound at replication foci, calculated as per the formulas for $E_{RPA1t}$ (1) and $F_{RPA1t}$ (2) for the photobleached and non-photobleached focus, respectively. (**e**) Normalized intensity of the photobleached RPA1 focus (as per formula 5).

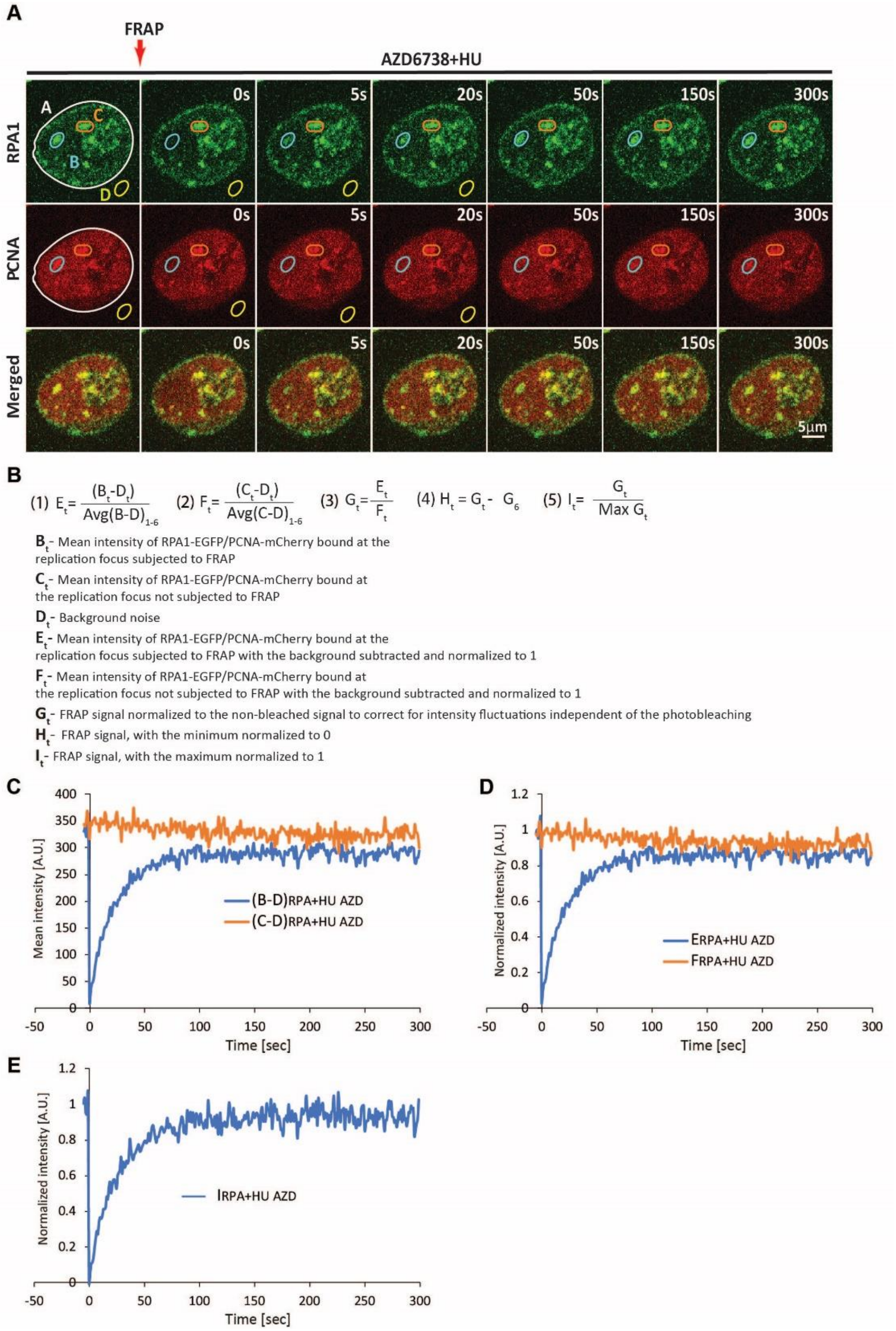


**Figure S5. Cell cycle progression following acute hydroxyurea-induced replication stress.** (**a**) Representative timelapse images of PCNA and RPA1 through 24-h after HU treatment and wash-out. (**b**) Same as (a), but under conditions of ATR inhibition (3μM AZD6738) before, during, and after HU treatment. (**c**) Same as (a) and (b), but under conditions of combined ATR+ATM (10μM Ku55933) inhibition. Scale bars = 5 μm. Timepoints of HU addition and washout are indicated by red arrows. (**d**) Progression of cells through the cell cycle following acute HU-induced replication stress under the above-described conditions. For HU: n = 22 cells; for HU+AZD: n = 44 cells; for HU+AZD+KU: n = 47 cells. **Abbreviations**: HU: hydroxyurea; AZD: AZD6738; KU: Ku55933

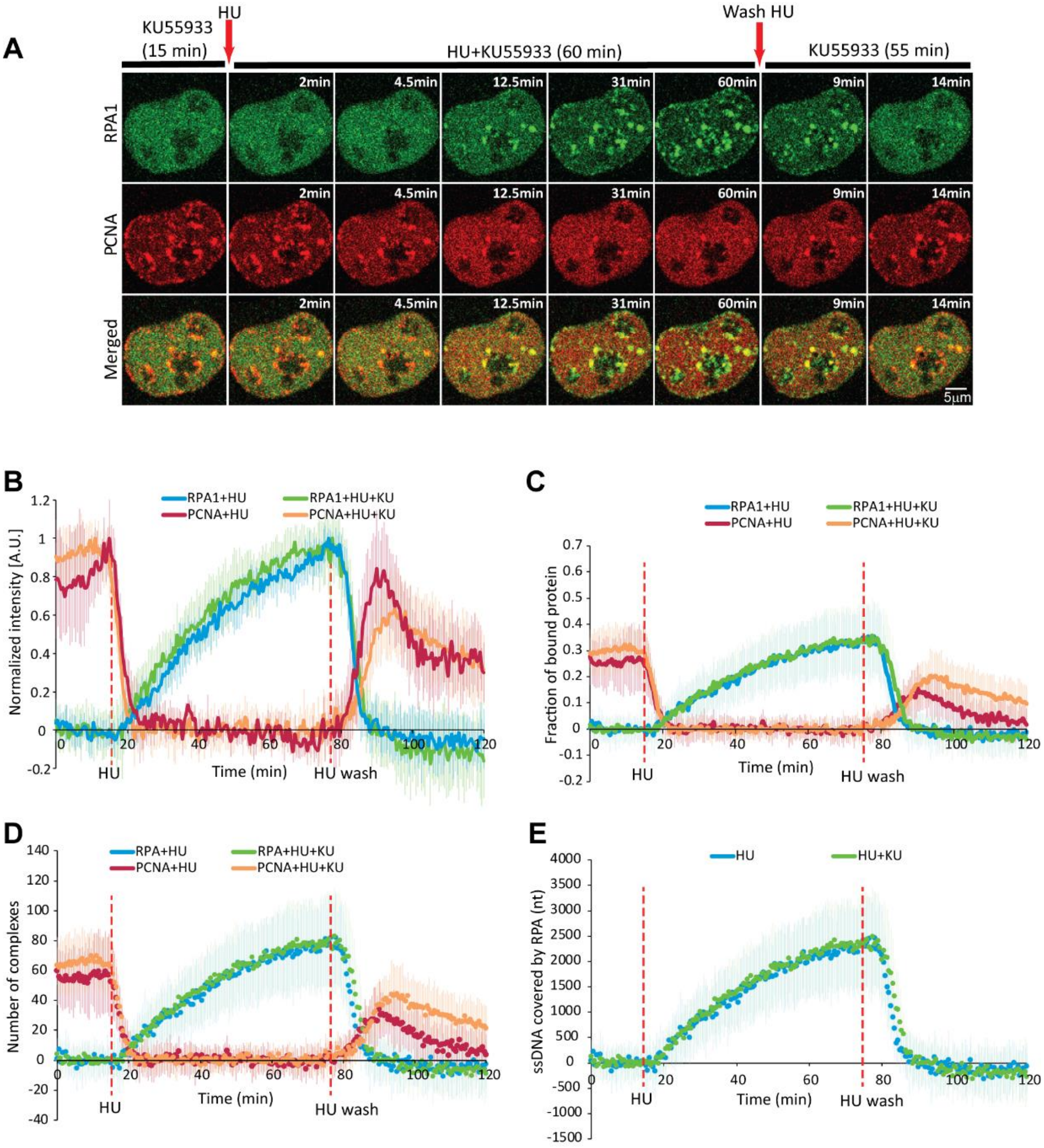


**Figure S6. Influence of ATM activity on PCNA and RPA1 dynamics during hydroxyurea-induced fork stalling and restart.** (**a**) Representative time-lapse images of RPA1 and PCNA before, during, and after HU treatment under ATM inhibition (10µM Ku55933). Arrows indicate timepoints of HU addition and washout. Scale bar = 5 µm. (**b**) Normalized kinetics of PCNA and RPA1 at replication foci during HU-induced replication fork stalling and restart, with or without ATM inhibition. The maximum intensity of PCNA/RPA1 at replication foci is normalized to 1. (**c**) Fraction of PCNA and RPA1 bound at replication foci during HU-induced replication fork stalling and restart, with or without ATM inhibition, relative to the total nuclear intensity of PCNA/RPA1, which is normalized to 1. (**d**) Estimated number of PCNA homotrimer and RPA heterotrimer complexes engaged at replication foci with or without ATR inhibition. (**e**) Average length of ssDNA (nt) covered by RPA at a single replication fork with versus without ATM inhibition. Dashed green lines indicate the timepoints of HU addition and washout. **Abbreviations**: HU: hydroxyurea; KU: Ku55933

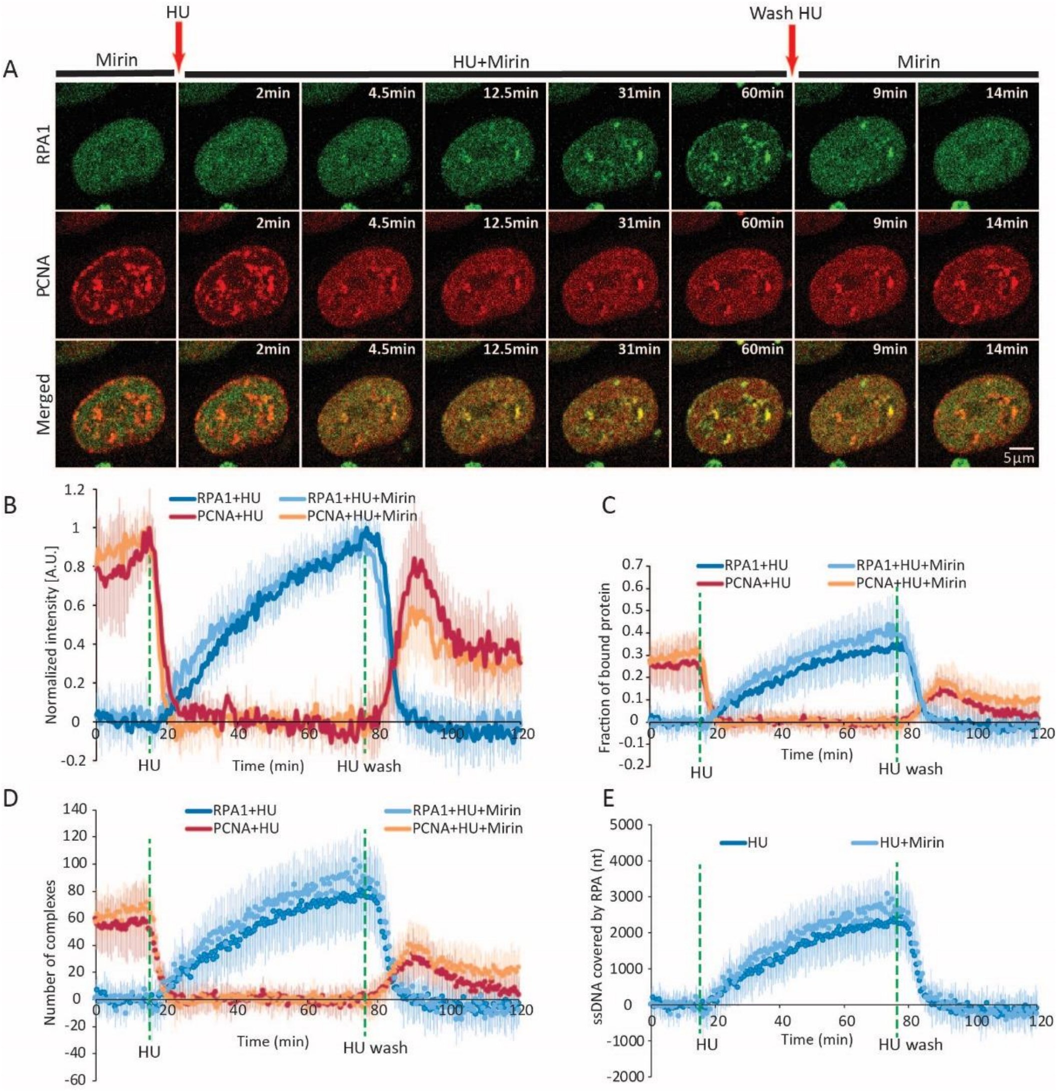


**Figure S7. Influence of MRE11 activity on PCNA and RPA1 dynamics during HU-induced replication fork stalling and restart.** (**a**) Representative time-lapse images of RPA1 and PCNA before, during, and after HU treatment under MRE11 inhibition (100μM mirin). Arrows indicate timepoints of HU addition and washout. Scale bar = 5 μm. (**b**) Normalized kinetics of PCNA and RPA1 recruited at replication factories during HU-induced replication fork stalling and restart, with or without MRE11 inhibition. The maximum intensity of PCNA/RPA1 engaged at replication factories is normalized to 1. (**c**) Fraction of PCNA and RPA1 bound at replication foci during HU-induced replication fork stalling and restart, with or without MRE11 co-inhibition, relative to the total nuclear intensity of PCNA/RPA1, which is normalized to 1. (**d**) Estimated number of PCNA homotrimer and RPA heterotrimer complexes engaged at replication foci, with or without MRE11 inhibition. (**e**) Estimated number of nucleotides covered by RPA heterotrimers with or without MRE11 inhibition. Dashed green lines indicate timepoints of HU addition and washout. Data presented as the mean±SD. For HU+Mirin: n = 10 cells; for HU: n = 17 cells. **Abbreviations**: HU: hydroxyurea

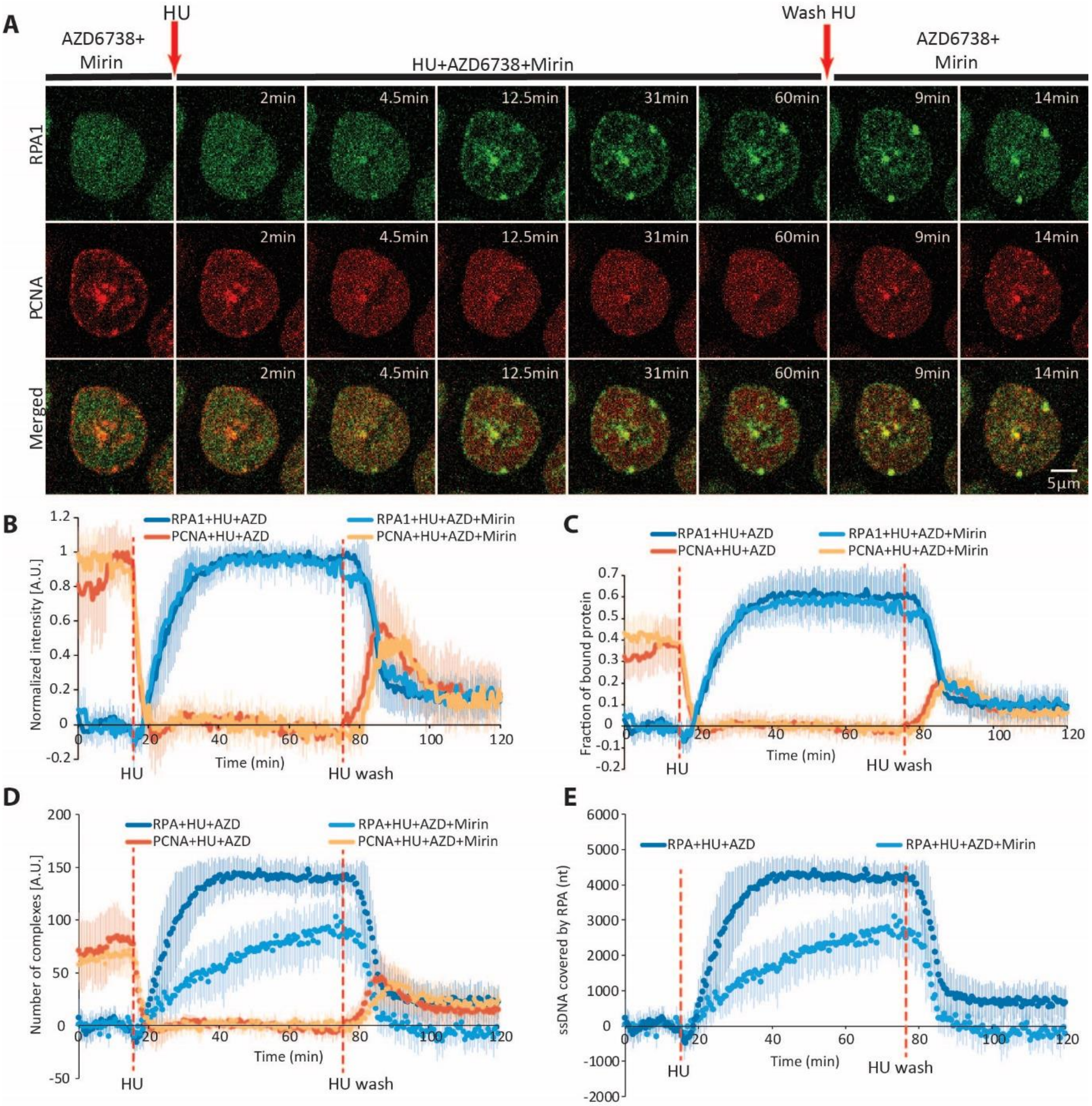


**Figure S8. Influence of MRE11 activity on PCNA and RPA1 dynamics during HU-induced replication fork stalling and restart under conditions of ATR inhibition.** (**a**) Representative time-lapse images of RPA1 and PCNA before, during, and after HU treatment under combined MRE11 (Mirin) and ATR (3μM AZD6738) inhibition. Arrows indicate timepoints of HU addition and washout. Scale bar = 5 μm. (**b**) Normalized kinetics of PCNA and RPA1 recruited at replication factories during HU-induced replication fork stalling and restart under ATR inhibition (3μM AZD6738) with or without MRE11 co-inhibition (mirin). The maximum intensity of PCNA/RPA1 engaged at replication foci is normalized to 1. (**c**) Fraction of PCNA and RPA1 bound at replication foci during HU-induced replication fork stalling and restart under ATR inhibition, with or without MRE11 co-inhibition, relative to the total nuclear intensity of PCNA/RPA1, which is normalized to 1. (**d**) Estimated number of PCNA homotrimer and RPA heterotrimer complexes engaged at replication foci under ATR inhibition, with or without MRE11 inhibition. (**e**) Estimated number of nucleotides covered by RPA heterotrimers with or without MRE11 inhibition. Dashed orange lines indicate the timepoints of HU addition and washout. For HU+mirin+AZD: n = 15 cells; for HU+mirin: n = 10 cells. **Abbreviations**: HU: hydroxyurea; AZD: AZD6738

**Supplementary Table 1.** Half-times of PCNA/RPA1 removal and loading during fork stalling and restart.

| | HU addition | | HU wash-out | |
|---|---|---|---|---|
| Treatment | PCNA removal $t_{1/2}$ (min) | RPA1 loading $t_{1/2}$ (min) | PCNA loading $t_{1/2}$ (min) | RPA1 removal $t_{1/2}$ (min) |
| - | 2.06±0.85 | 23.91±2.12 | 5.09±1.98 | 3.24±1.19 |
| ATRi | 1.70±0.48 | 5.60±2.21 | 4.40±0.93 | 2.75±1.43 |
| ATMi | 1.97±0.66 | 20.16±5.59 | 5.71±5.23 | 3.89±1.65 |
| ATRi+ATMi | 2.15±0.78 | 4.35±1.25 | 3.60±1.33 | 4.10±1.43 |
| MRE11i | 1.81±0.39 | 21.00±2.04 | 5.85±2.98 | 2.70±0.82 |
| ATRi+MRE11i | 1.60±0.60 | 5.63±2.06 | 6.07±2.09 | 2.63±1.36 |

**Supplementary Table 2.** Number of complexes engaged per replication fork during stalling.

| Treatment | PCNA before HU | RPA loaded after HU addition | ssDNA (nt) after HU addition | PCNA after HU wash-out | Residual RPA after HU wash-out |
|---|---|---|---|---|---|
| - | 59.01±22.6 | 80.71±27.74 | 2421±832 | 33.69±16.6 | - |
| ATRi | 66.94±24.30 | 139.33±13.76 | 4180±413 | 42.71±22.26 | 22.23±16.03 |
| ATMi | 65.77±18.83 | 77.67±30.9 | 2330±926 | 41.54±18.61 | - |
| ATRi+ATMi | 69.64±25.13 | 141.32±23.06 | 4240±692 | 75.95±26.23 | 74.06±18.63 |
| MRE11i | 65.46±17.37 | 89.48±25.11 | 2685±753 | 40.51±17.45 | - |
| ATRi+MRE11i | 89.54±23.84 | 134.28±25.88 | 4028±776 | 41.98±19.83 | 23.7±17.07 |

**Supplementary Table 3.** Rate (in complexes) of PCNA/RPA1 removal and loading during for stalling and restart.

| | HU addition | | HU wash-out | |
|---|---|---|---|---|
| **Treatment** | **PCNA removal (complexes/min)** | **RPA loading (complexes/min)** | **PCNA loading (complexes/min)** | **RPA removal (complexes/min)** |
| - | 14.95±6.94 | 1.33±0.45 | 2.83±2.49 | 8.91±5.2 |
| ATRi | 14.99±8.22 | 9.61±3.57 | 4.87±1.88 | 11.59±6.09 |
| ATMi | 11.64±6.82 | 1.31±0.42 | 3.37±1.53 | 8.48±4.1 |
| ATRi+ATMi | 14.10±5.89 | 10.17±3.72 | 10.28±4.79 | 7.15±3.51 |
| MRE11i | 13.33±5.79 | 1.99±0.67 | 3.93±2.47 | 16.96±3.84 |
| ATRi+MRE11i | 17.99±6.52 | 9.89±3.22 | 8.73±2.31 | 13.25±7.45 |

**Supplementary Table 4.** Exchange rate of complexes at replication forks during stalling and restart.

| | **Treatment** | **$t_{1/2}$ exchange (s)** | **Exchange rate (complexes/min)** |
|---|---|---|---|
| | - | 55±6.25 | 31.2±4.9* |
| **PCNA** | ATRi | 58±10.2 | 39.41±6.09* |
| | HU | 149±28.35 | 2.38±0.91* |
| | HU+ATRi | 153±34.35 | 2.71±0.99* |
| **RPA1** | HU | 5±0.95 | 407.45±94.82** |
| | HU+ATRi | 29±4.26 | 228.12±65.67** |

**exchange rate measured during the first 1 min of recovery after photobleaching*

***exchange rate measured during the first 9 sec of recovery after photobleaching*
